# A peptidomimetic inhibitor of *Vibrio cholerae* adhesin protein FrhA

**DOI:** 10.64898/2026.09.07.749730

**Authors:** Mingyu Wang, Charity Yongo-Luwawa, Laura Zhang, Ryu Williston, John Presley, Zhaleh Mehboodi, Dieter P. Reinhardt, Karl E. Klose, William D. Lubell, Shuaiqi Guo

**Author notes:** Equal contributions (Mingyu Wang, Charity Yongo-Luwawa, Laura Zhang).

## Abstract

Cholera is a diarrheal disease resulting from colonization of the small intestine by the Gram-negative bacterium *Vibrio cholerae*. The peptide-binding domain (PBD) of the adhesion protein FrhA is critical for *V. cholerae* intestinal colonization and biofilm formation. Antagonists of the FrhA-PBD were designed based on the tripeptide motif H-Trp-Thr-Asp-OH. Constraining the N-terminal tryptophan as a 1,2,3,4-tetrahydro-β-carboline-3-carboxylic acid (Tcc) residue and X-ray analysis of a co-crystal with the FrhA-PBD define the minimal inhibitory motif. Moreover, H-(R)-Tcc-Thr-Asp-OH exhibited nanomolar binding affinity (Kd = 101 ± 30.2 nM) to the FrhA-PBD, blocking adhesion protein-mediated bacterial hemagglutination and reducing biofilm formation *in vitro*. Intestinal epithelial cell adhesion to the FrhA-PBD was also blocked by the Tcc tripeptide (IC□□ = 24 µM; 95% CI, 11–51 µM), which resisted chymotrypsin proteolysis. The activity of this peptidomimetic inhibitor and the atomic-level understanding of FrhA-PBD binding provide insights for developing anti-adhesion therapies to treat cholera.

## INTRODUCTION

Cholera is a diarrheal disease and persistent public health threat caused by the waterborne pathogen *Vibrio (V.) cholerae*^1–4^. After ingestion of contaminated food or water, the Gram-negative bacterium adheres to the small intestine, where colonization promotes pathogenesis^2, 5^. Upon small intestine colonization, *V. cholerae* releases cholera toxin, which can activate the stimulatory alpha subunit in the heterotrimeric G protein to initiate a cAMP-dependent signalling pathway leading to stimulation of chloride efflux with consequent osmotic movement of water into the intestinal lumen and diarrhea. ^6–9^ Standard treatments for cholera infection, including oral rehydration therapy, antibiotics, and intravenous fluid replacement for severe cases, have limitations in availability and effectiveness^3^. With the emergence of antibiotic-resistant *V. cholerae* strains exacerbated by climate change,^10–14^ interest mounts in developing novel therapeutic strategies to combat infection. In this context, anti-adhesion strategies offer promise for reducing bacterial virulence by limiting host colonization without directly inhibiting bacterial growth or imposing selective pressure for antimicrobial resistance^15–23^.

The potential of anti-adhesion therapy is reinforced by the entry into phase 1b human clinical trials of GSK3882347, an orally administered competitive antagonist of the *Escherichia coli* adhesin FimH for the treatment of uncomplicated urinary tract infection^24–26^. Furthermore, the small molecule C607-0736 is a competitive inhibitor of the virulence and resistance factor OmpV, which enhances *V. cholerae* adhesion and internalization^27^. The colonization of *V. cholerae* in the infant mouse intestine was blocked by C607-0736, which also reduced toxin levels in host cells^27^.

The Flagellar Regulated Hemagglutinin A (FrhA) adhesin protein of *V. cholerae* is critical for intestinal colonization, human epithelial cell binding, hemagglutination, and biofilm formation^1, 2, 28–34^. The elongated modular architecture of FrhA promotes attachment of *V. cholerae* to host surfaces^35–37^. Anchored on the bacterial outer membrane, FrhA engages host targets with multiple functional domains, including a peptide-binding domain (PBD) implicated in intestinal attachment^37–39^.

Towards the validation of the FrhA-PBD as an anti-adhesion target, the L-pentapeptide FrhA-PBD ligand H-Ala-Gly-Tyr-Thr-Asp-OH (L-**1**) was shown to weakly inhibit *V. cholerae* colonization of the mouse intestine^29, 40^. Moreover, X-ray analyses have been obtained of L-**1** and L-**2** (H-Ala-Gly-Trp-Thr-Asp-OH) in complex with a recombinant protein construct, FrhA_Split-PBD_, containing the PBD and its adjacent “Split” domain^41^. In the X-ray structure, coordination of the *C*-terminal carboxylate to two calcium ions in the shallow binding pocket of FrhA-PBD anchors peptide L-**1**, which also forms multiple hydrogen bonds with the protein^29^. A tyrosine residue on the protein surface near the binding pocket has been suggested to offer potential for engaging the ligand through hydrophobic interactions^29, 41^. Subsequently, the truncated motifs H-Tyr-Thr-Asp-OH (L-**3**) and H-Trp-Thr-Asp-OH (L-**4**) were shown to exhibit higher binding affinity than pentapeptide L-**1**^41^. Proteolytic degradation in the protease-rich intestine may account partly for the weak activity of the L-peptide FrhA-PBD ligands in the mouse colonization model^16^.

To improve binding affinity and reduce metabolic susceptibility, noncanonical and conformationally constrained amino acids are being examined in next generation adhesin antagonists^42–45^. Attempts herein to replace the penultimate residue in the tripeptide with a set of threonine surrogates in analogs **5**-**10** resulted however, in significant losses of affinity. On the other hand, inversion of the configuration of the *N*-terminal aromatic residue provided respectively D-tyrosine and D-tryptophan inhibitors D-**3** and D-**4**, which exhibited comparable if not better binding affinity than the L-counterparts L-**3** and L-**4**.

The 1,2,3,4-tetrahydro-β-carboline-3-carboxylate (Tcc) residue has now been explored as a conformationally rigid analog of tryptophan, which constrains the indole side chain and backbone amine within a tricyclic scaffold.^46^. Introducing this rigid residue into the H-Trp-Thr-Asp-OH motif provided H-(*S*)-Tcc-Thr-Asp-OH (*S*-**11**), which was crystallized in complex with FrhA-PBD. Furthermore, employing the constrained amino acid diastereomer, H-(*R*)-Tcc-Thr-Asp-OH (*R*-**11**) was synthesized and shown to bind the adhesin protein with a dissociation constant (K_d_) in the nanomolar range. The functional potency of D-Trp and (*R*)-Tcc inhibitors was examined in hemagglutination^29, 47, 48^, biofilm formation^29^, and human intestinal epithelial cell adhesion assays. In addition, the influences of peptide length and the configuration and structure of the aromatic residue on proteolytic susceptibility were assessed using chymotrypsin as a major intestinal protease^49–51^.

## RESULTS

### Chemistry

A panel of penta- and tripeptide analogues was designed to study structure-activity relationships with respect to binding affinity, as well as effects on hemagglutination, biofilm formation, intestinal cell adhesion and proteolytic stability (Table 1). Considering earlier studies^40, 41^, stereochemical inversions and conformational restriction were explored primarily at the aromatic and threonine residues. Notably, restraint of the tryptophan χ dihedral angle was explored employing the Tcc residue, which respectively favours gauche (−) and (+) conformers contingent on residue location at the *N*-terminal and central positions of the peptide sequence (Figure 1)^52–54,55^. Employing the Pictet-Spengler reaction on 2-chlorotrityl chloride resin^46^, Tcc tripeptides *R*- and *S*-**11** were synthesized using pyridinium p-toluenesulfonate to catalyze cyclization without removal of acid labile groups^56^. Upon cleavage of resin **13** using 20% HFIP in dichloromethane (DCM)^57^, Pictet-Spengler reaction conversion was evaluated by liquid chromatography-mass spectrometry (LC-MS) analysis of protected Tcc peptide **14**. Resin cleavage and removal of side chain protection were consequently performed using trifluoroacetic acid (TFA) in DCM to obtain tripeptides *R*-and *S*-**11** of >95% purity (Scheme 1 and Table S1).

**Figure 1.**
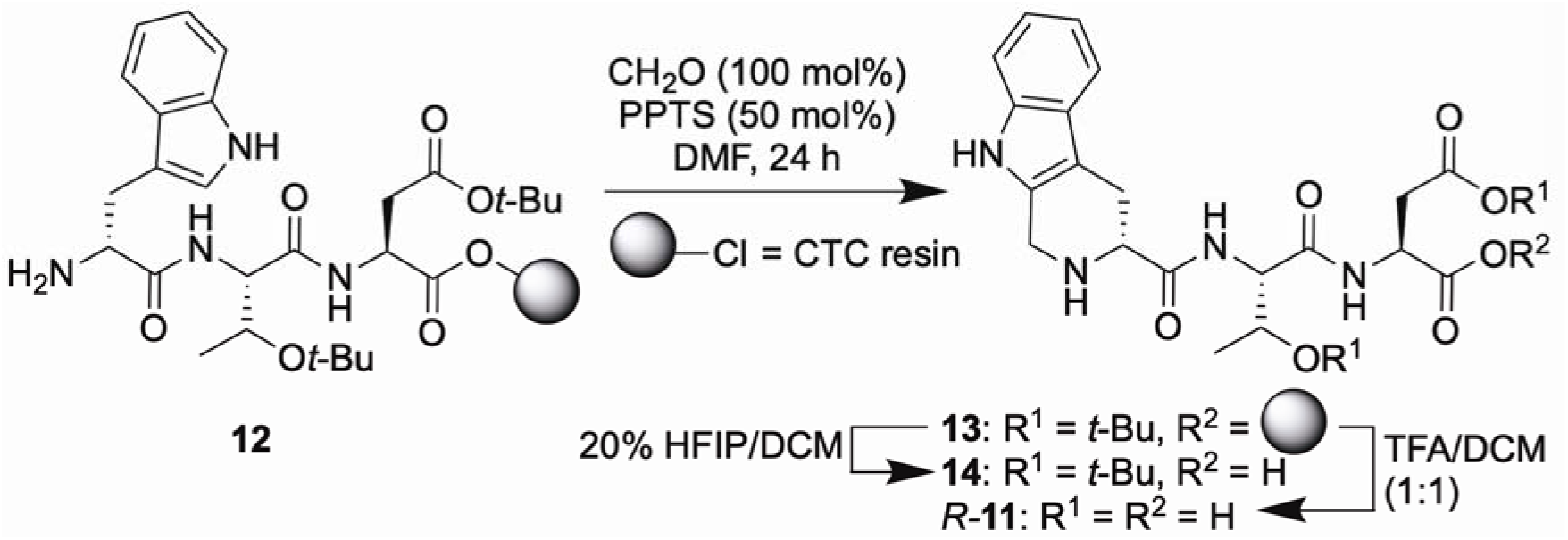
Tcc residue in gauche (-) and (+) conformers.

**Scheme 1.**
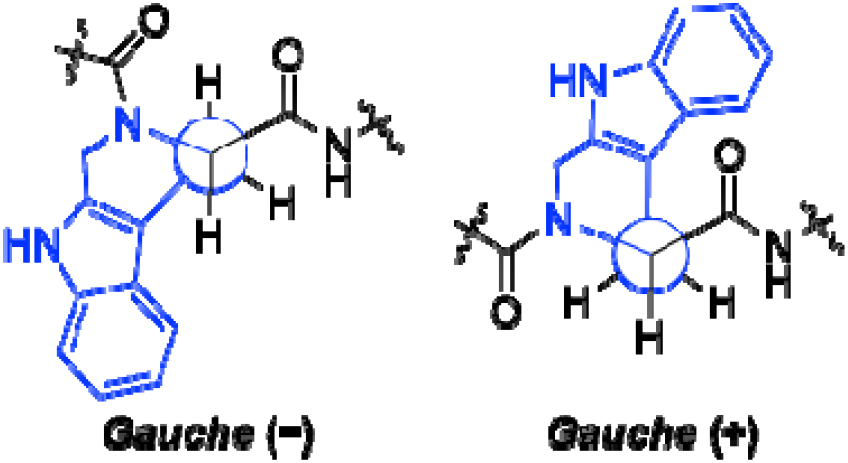
Synthesis of H-(*R*)-Tcc-Thr-Asp-OH (*R*-**11**).

### Binding affinity and biological assessments

The amino acid sequence of the tripeptide H-Tyr-Thr-Asp-OH (L-**3**) was previously identified as a minimal motif required for binding to the FrhA-PBD^41^. In the current study, the *C*-terminal aspartate residue was not examined in part due to its importance in engaging calcium ions in the binding pocket as observed in the X-ray structure of pentapeptide L-**1** in complex with the FrhA-PBD. Furthermore, substitution of natural L-Asp with D-Asp in pentapeptide L-**1** reduced binding affinity approximately sixfold, increasing the dissociation constant from 500 nM to 3.0 µM^41^. Likewise, substitution of the *C*-terminal carboxylic acid with a carboxamide counterpart abolished binding to the homologous conserved peptide-binding domain of the *Marinomonas primoryensis* ice-binding protein (*Mp*IBP-PBD)^40^.

**Table 1.** FrhA-PBD inhibitor binding affinity, bioactivity, and chymotrypsin stability.

| # | Compound name | MST $K_d \pm SD$ (nM) | HA IC<sub>50</sub> ± SD (μM) | Biofilm IC<sub>50</sub>, μM (95% CI) | Caco-2 IC<sub>50</sub>, μM (95% CI) | Chymotrypsin digest half life (min) |
| --- | --- | --- | --- | --- | --- | --- |
| L-1 | H-Ala-Gly-Tyr-Thr-Asp-OH | 500 ± 140<sup>41</sup> | >100 ref | 13 (8.3–17) | 220 (130–400) | $t_{1/2}$ = ~58.1 |
| L-2 | H-Ala-Gly-Trp-Thr-Asp-OH | 380 ± 100<sup>41</sup> | — | — | — | $t_{1/2}$ = 20.3 ± 4.4 min |
| D-2 | H-Ala-Gly-D-Trp-Thr-Asp-OH | — | — | — | — | N.D. |
| L-3 | H-Tyr-Thr-Asp-OH | 191 ± 44.0 | — | — | — | N.D. |
| D-3 | H-D-Tyr-Thr-Asp-OH | 69.3 ± 19.8 | — | — | — | N.D. |
| L-4 | H-Trp-Thr-Asp-OH | 244 ± 116 | 106 ± 14.1 | — | — | N.D. |
| D-4 | H-D-Trp-Thr-Asp-OH | 110 ± 43.5 | 11.1 ± 1.06 | 1.6 (0.80–2.5) | 44 (31–63) | N.D. |
| 5 | H-Tyr-D-Thr-Asp-OH | 3,040 ± 1,110 | — | — | — | — |
| 6 | H-Tyr-allo-Thr-Asp-OH | 7,380 ± 2,330 | — | — | — | — |
| 7 | H-Tyr-D-allo-Thr-Asp-OH | 32,400 ± 14,300 | — | — | — | — |
| 8 | H-D-Tyr-allo-Thr-Asp-OH | 1,460 ± 477 | — | — | — | — |
| 9 | H-D-Tyr-D-Thr-Asp-OH | 4,630 ± 1,830 | — | — | — | — |
| 10 | H-D-Tyr-Hyp-Asp-OH | N.B. | — | — | — | — |
| S-11 | H(S)-Tcc-Thr-Asp-OH | 374 ± 153 | 192 ± 39.5 | — | 210 (150–290) | N.D. |
| R-11 | H(R)-Tcc-Thr-Asp-OH | 101 ± 30.2 | 27.9 ± 6.64 | 2.2 (0.95–3.4) | 24 (11–51) | N.D. |

MST: Microscale thermophoresis; HA: hemagglutination; 95% CI: 95% confidence interval; N.B.: no binding detected; N.D.: Negligible digestion detected after 120 min.

Stereoisomeric substitution was explored at the neighboring L-threonine residue. Effects on binding affinity were examined using microscale thermophoresis (MST)^41^. Replacement of L-threonine by D-threonine, L-allothreonine, and D-allothreonine in H-Tyr-Thr-Asp-OH (L-**3**, K_d_ = 191 ± 44.0 nM)^41^ caused 16- to 170-fold decreases in binding affinity (Figure 2). Similarly, replacement of L-threonine by D-threonine and L-allothreonine in the more potent H-D-Tyr-Thr-Asp-OH (**D-3**, K_d_ = 69.3 ± 19.8 nM)^41^ caused 67- and 21-fold reductions, respectively, in binding affinity (Figure 2). Moreover, replacement of L-threonine with L-hydroxyproline (Hyp) as a constrained counterpart gave H-D-Tyr-Hyp-Asp-OH (**10**), which showed no detectable binding affinity in the MST assay. In crystal structures of pentapeptide L-**1** (PDB ID: 9YBQ) and L-Trp counterpart H-Ala-Gly-Trp-Thr-Asp-OH (L-**2**; PDB ID: 9Y9W) in complex with FrhA-PBD, the backbone and side chain of L-threonine make hydrogen bonds with the binding pocket which are likely disrupted on modifications of stereochemistry and structure in tripeptides L-**3** and D-**3**, leading to reduced binding affinity.

**Figure 2.**
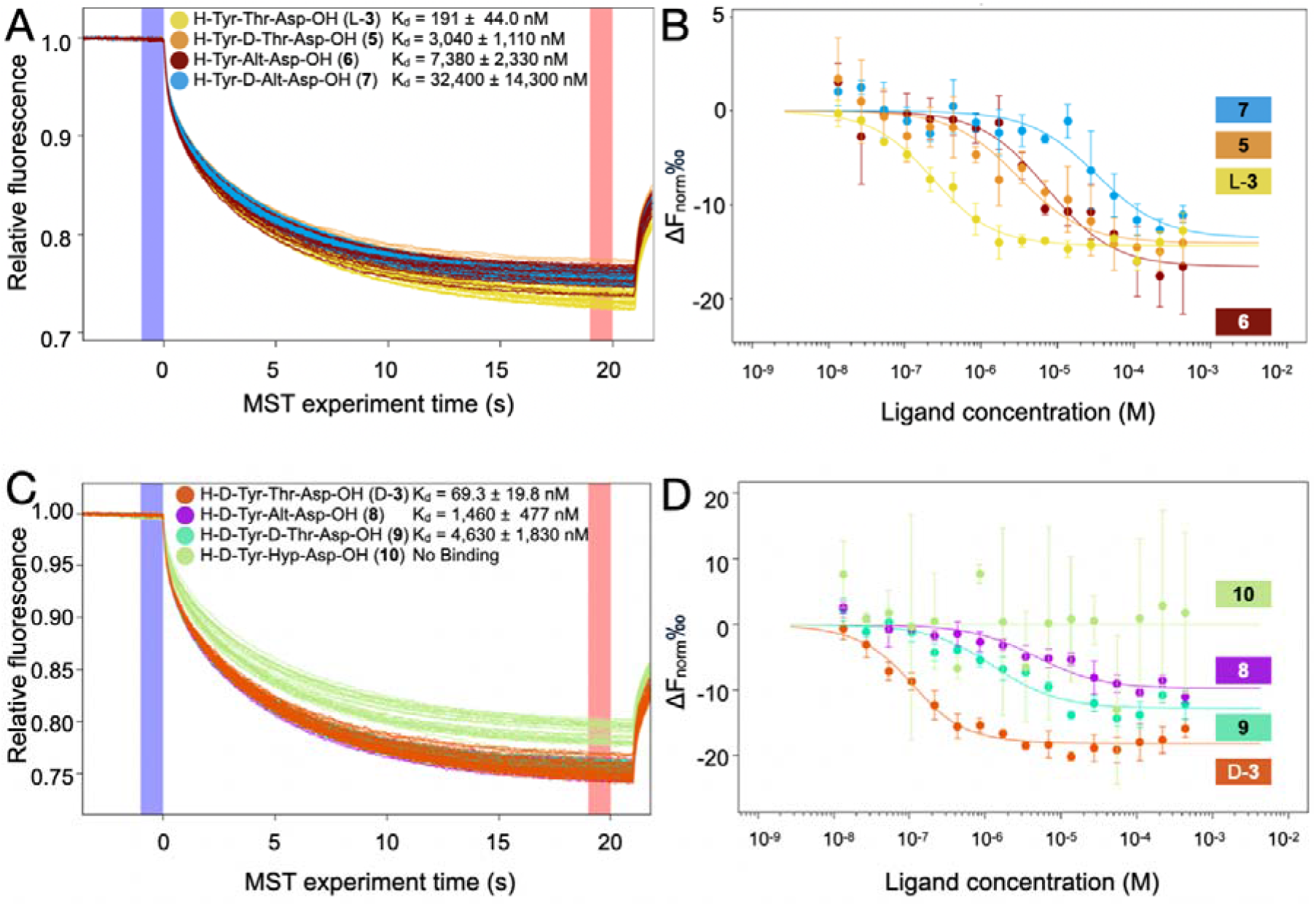
Modification of Thr in H-D- and L-Tyr-Thr-Asp-OH (D- and L-3) reduces binding affinity to FrhA_Split-PBD_. (Left) Representative MST traces and (right) corresponding binding curves for FrhA_Split-PBD_ with peptide analogues **3, 5**-**10**. Binding curves were generated by plotting the change in normalized fluorescence (ΔF_norm_) as a function of ligand concentration. Experiments were performed in triplicate; fitted K_d_ values are indicated in the figure. Lowercase amino acid abbreviations denote D-configured residues; Alt and D-Alt denote L-allothreonine and D-allothreonine, respectively.

At the *N*-terminal aromatic residue, D- and L-tyrosine were replaced with the corresponding tryptophan residues in H-D- and L-Trp-Thr-Asp-OH (D-**4** and L-**4**) which exhibited K_d_ values of 110 ± 43.5 nM and 244 ± 116 nM, respectively^41^. Employing on-resin Pictet-Spengler chemistry^46^, the corresponding (*R*)- and (*S*)-Tcc counterparts were synthesized to restrain the aromatic residue side chain to gauche c-space. Compared with the tripeptides D**-4** and L**-4**, constrained analogues H-(*R*)- and (*S*)-Tcc-Thr-Asp-OH exhibited, respectively, similar and 1.5-fold weaker binding affinities: K_d_ = 101 ± 30.2 nM and 374 ± 153 nM (Figure 3).

**Figure 3.**
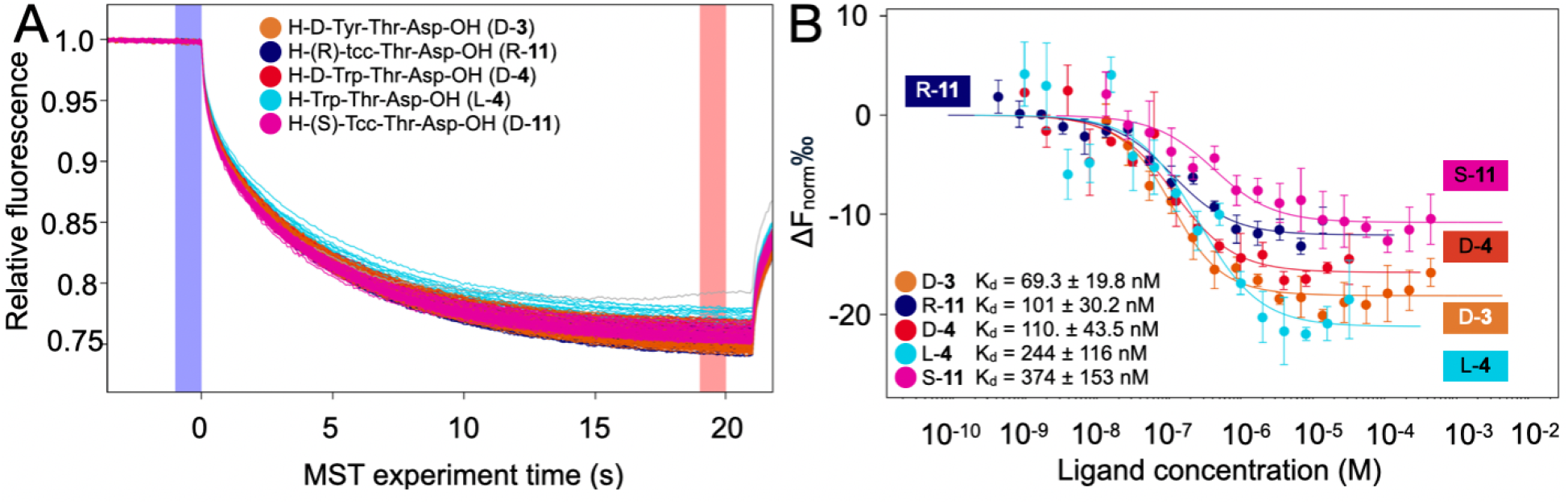
Influence of aromatic residue on binding affinity: (left) representative MST traces and (right) corresponding binding curves. Fitted K_d_ values are indicated in the figure.

### X-ray Crystal Structure Analysis of FrhA_Split-PBD_ in complex with (*S*)-Tcc-Thr-Asp-OH

The X-ray co-crystal structure of H-(*S*)-Tcc-Thr-Asp-OH (*S*-**11**) was obtained in complex with the recombinant protein construct FrhA_Split-PBD_ (Figure 4, Figure S1, Table 2, and Table S2). In this first structure of a truncated inhibitor bound to FrhA_Split-PBD_, the threonine and aspartic acid residues interact respectively with the peptide-binding pocket and Ca^2^□ ions in a manner similar to that observed for pentapeptide L-**2** bound to the same protein (PDB ID: 9Y9W).³□ Like the tryptophan side chain of pentapeptide L-**2**, the aromatic moiety of (*S*)-Tcc points away from the binding pocket into the solvent.

**Figure 4.**
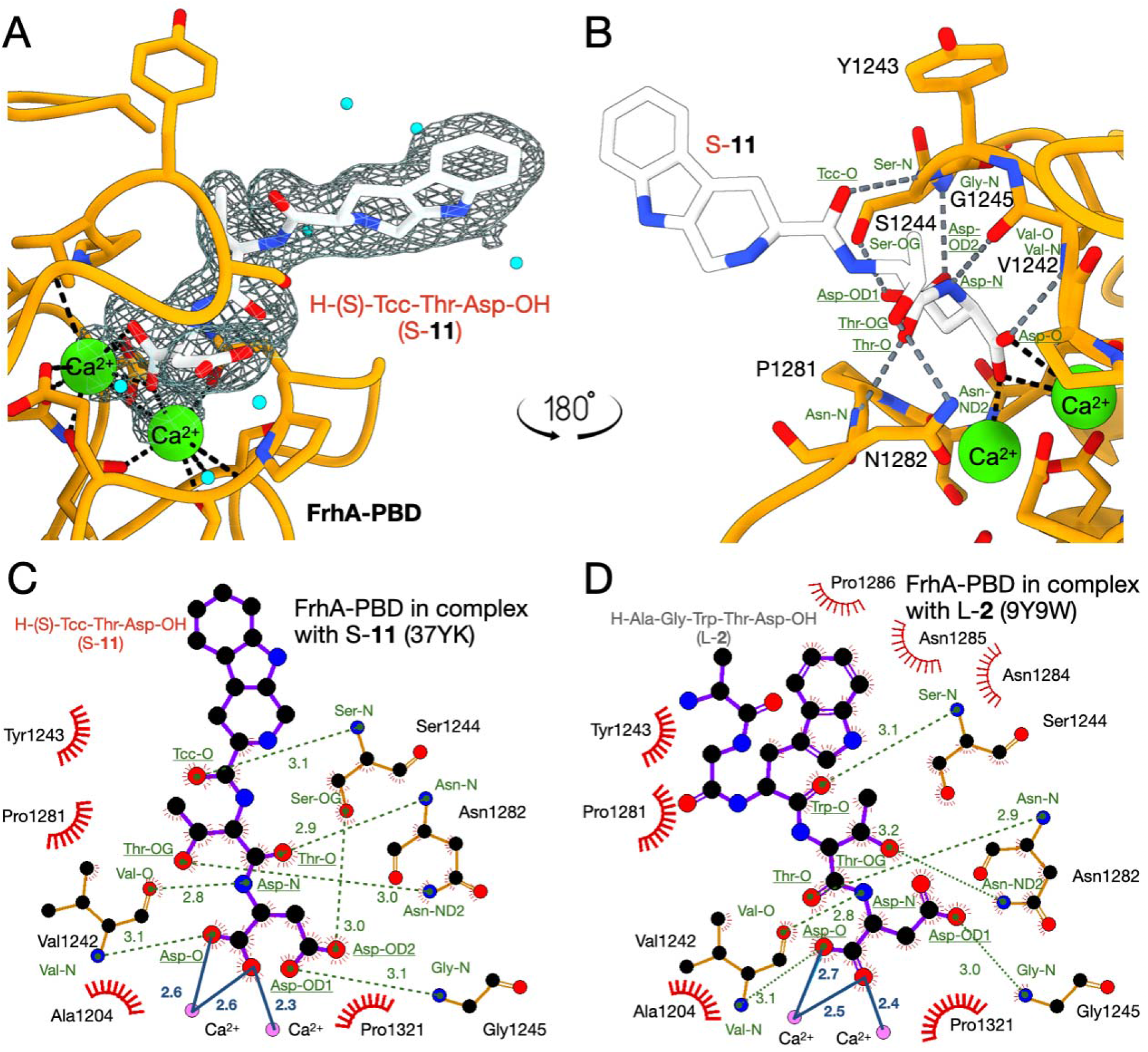
X-ray crystal structure of FrhA_Split-PBD_ in complex with H-(*S*)-Tcc-Thr-Asp-OH (*S*-11) at 2.2-Å resolution. Electron density (top left) surrounding bound *S*-**11** (white) as a mesh, with the two Ca^2^o ions in the ligand-binding site shown as green spheres and water molecules as cyan spheres. Bound *S*-**11** (top right) with hydrogen bonds (grey dashed lines) and ligand–Ca^2^o coordination (black). LigPlot diagrams of the binding interactions of *S*-**11** (bottom left) and pentapeptide L-**2** (bottom right), respectively, with surrounding FrhA-PBD residues and coordinating Ca^2^o ions. Green dashed lines indicate hydrogen bonds, and solid blue lines indicate ligand–Ca^2^o coordination. Distances are given in Å. For L-**2**, thinner green dashed lines indicate hydrogen bonds observed in at least one, but not all, of the six chains in the asymmetric unit (Table 2).

**Table 2.** Calcium coordination and hydrogen bonds at the FrhA_Split-PBD_–ligand interface.

| FrhA<sub>Split-PBD</sub> atom | H-(S)-Tcc-Thr-Asp-OH (S-11, PDB ID: 37YK) | H-Ala-Gly-Trp-Thr-Asp-OH (L-2, PDB ID: 9Y9W) | H-Ala-Gly-Tyr-Thr-Asp-OH (L-1, PDB ID: 9YBQ) |
| --- | --- | --- | --- |
| $\text{Ca}^{2+}$ coordination to the Asp α-carboxylate | | | |
| $\text{Ca}^{2+}$ | Asp-α-carboxylate O (Asp-O), 2.3 Å | Asp-O, 2.6 Å | Asp-O, 2.4 Å |
| $\text{Ca}^{2+}$ | Asp-O, 2.6 Å | Asp-O, 2.5 Å | Asp-O, 2.5 Å |
| $\text{Ca}^{2+}$ | Asp-O, 2.6 Å | Asp-O, 2.6 Å | Asp-O, 2.7 Å |
| N-terminal aromatic residue |  |  |  |
| Ser1244-N | Tcc-O, 3.1 Å | Trp-O, $3.1 \pm 0.1$ Å | Tyr-O, $3.2 \pm 0.1$ Å (5/6)* |
| Thr residue |  |  |  |
| Asn1282-N | Thr-O, 2.9 Å | Thr-O, $2.9 \pm 0.1$ Å | Thr-O, $3.0 \pm 0.1$ Å |
| Asn1282–ND2 | Thr–OG, 3.0 Å | Thr–OG, 3.2 ± 0.1 Å (5/6)* | — |
| C-terminal Asp residue |  |  |  |
| Val1242–O | Asp–N, 2.8 Å | Asp–N, 2.8 ± 0.0 Å | Asp–N, 2.8 ± 0.1 Å |
| Val1242–N | Asp–O, 3.1 Å | Asp–O, 3.1 ± 0.1 Å (4/6)* | Asp–O, 3.2 ± 0.2 Å |
| Gly1245–N | Asp–OD, 3.1 Å | Asp–OD, 3.0 ± 0.2 Å (5/6)* | Asp–OD, 3.2 ± 0.1 Å (2/6)* |
| Ser1244–OG | Asp–OD, 3.0 Å | — | Asp–OD, 3.0 ± 0.1 Å (4/6)* |

\* Fractions in parentheses indicate the number of chains in which the corresponding hydrogen bond was observed, out of six independent crystals.

Binding of tripeptide *S*-**11** is stabilized by coordination to the Ca^2^□ ions and a network of hydrogen bonds with the protein. The calcium ions appear at similar distances (2.3, 2.6, and 2.6 Å) from the coordinating oxygen atoms of the aspartate α-carboxylate in tripeptide *S*-**11** compared to that of pentapeptides L-**1** and L-**2** in related X-ray structures (Table 2). The hydrogen-bond network between the PBD and tripeptide *S*-**11** includes the Asp a- and b-carboxylate oxygens (Asp-O, OD1 and OD2) accepting hydrogen bonds respectively from the backbone amide nitrogen of Val1242 (3.14 Å) and Gly1245 (3.07 Å) and the side-chain hydroxyl group of Ser1244 (3.03 Å). The Asp backbone NH also hydrogen bonds to the backbone carbonyl oxygen of Val1242 (2.83 Å). The Thr backbone carbonyl and side-chain hydroxyl oxygen share respectively hydrogen bonds with the backbone and side-chain amide nitrogen of Asn1282 (N, 2.87 Å and ND2, 3.02 Å). Finally, the Tcc carbonyl oxygen accepts a hydrogen bond from the backbone amide nitrogen of Ser1244 (3.06 Å). A similar set of generally longer hydrogen bonds appear in the X-ray structures of the pentapeptides L-**1** and L-**2** bound to FrhA-PBD (Figure 4, Table 2). For example, the Trp carbonyl oxygen of pentapeptide L-**2** shares a hydrogen bond with the backbone amide nitrogen of Ser1244 (3.1 ± 0.1 Å; 6/6 chains). Moreover, pentapeptides L-**1** and L-**2** lack consistency in certain hydrogen bonds across all chains in the independent crystals (Table 2 and Figure S1). The more consistent bonds between the PDB and tripeptide *S*-**11** compared to pentapeptides L-**1** and L-**2** correlate with the tighter binding affinity of the former.

### Inhibition of Bacterial Hemagglutination

Promising inhibitors exhibiting binding affinity to the FrhA-PBD were next examined for anti-adhesion activity in a set of functional assays. The hemagglutination assay has historically been used with *V. cholerae* to assess anti-adhesion activity by measuring inhibition of bacteria-mediated red blood cell agglutination^47, 58, 59^. Previously, the FrhA-PBD was implicated in the cross-linking of erythrocytes by wild-type (WT) *V. cholerae*, and genetic deletion of the domain in the Δ*frha-pbd* mutant abolished ability to cause hemagglutination^28, 29^. Furthermore, pentapeptide L-**1** was shown to block hemagglutination at approximately 100 µM likely by targeting the FrhA-PBD^29^.

Herein, the peptidomimetics H-(*R*)- and (*S*)-Tcc-Thr-Asp-OH (*R*-**11** and *S*-**11**) were benchmarked against tryptophan counterparts H-D- and L-Trp-Thr-Asp-OH (D-**4** and L-**4**) in blocking WT *V. cholerae* hemagglutination of type O erythrocytes (Figure 5, Figure S2, and Table S3). RBCs alone and incubated with the *V. cholerae* Δ*frha-pbd* mutant lacking the PDB of FrhA, both exhibited no hemagglutination. In contrast, complete hemagglutination occurred with WT *V. cholerae* after two hours. Inhibitors D- and L-**4** and *R*- and *S*-**11** were evaluated over a range of final concentrations from 1.0 mM to 7.8 µM. Consistent with the binding affinity studies, inhibitors D-**4** and *R*-**11** (IC_50_ = 11.1 µM and 27.9 µM, respectively) having *R* configuration at the *N*-terminal aromatic residue blocked hemagglutination with greater potency than the *S*-counterparts L-**4** and *S*-**11** (IC_50_ = 106 µM and 192 µM).

**Figure 5.**
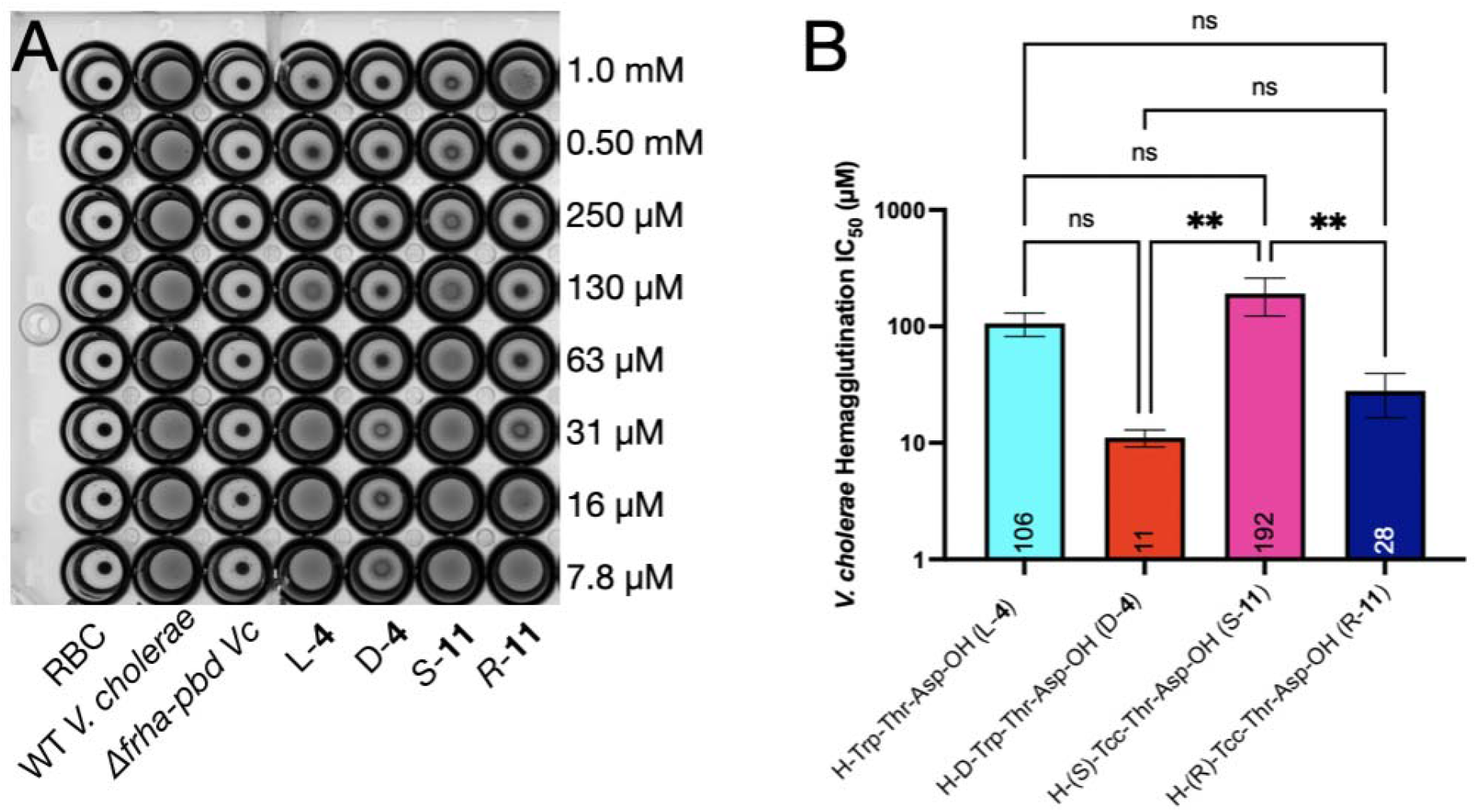
FrhA-PBD inhibitors block *Vibrio cholerae*-mediated hemagglutination. (Left) Representative experiments performed in triplicate of hemagglutination of human type O erythrocytes (RBC) and WT *V. cholerae* O395 in the presence of inhibitors over a twofold serial dilution from 1.0 mM to 7.8 mM, with controls of RBC alone and in the presence of WT and Δ*frha-pbd V. cholerae*. Plates were imaged after two hours at room temperature. (Right) Inhibitor IC_50_ values calculated quantitatively from dose–response curves of percentage hemagglutination inhibition over three separate experiments. Error bars show standard deviation (Table 1). See experimental section and supplementary information for details. One-way ANOVA with Tukey’s post-hoc test is indicated above the corresponding groups (Table S3); ns, not significant; P < 0.05 (*), P < 0.01 (**), P < 0.001 (***), and P < 0.0001 (****).

### Inhibition of Biofilm Formation by *V. cholerae*

The adhesion protein FrhA contributes significantly to *V. cholerae* biofilm formation, which is appreciably reduced relative to wild type in colonies of the Δ*frha-pbd* deletion mutant^28, 29^. Previously, pentapeptide L-**1** was shown to inhibit *V. cholerae* biofilm formation at 100 µM, likely by targeting the FrhA-PBD^29^. The relative potency against WT *V. cholerae* biofilm inhibition was compared using pentapeptide L-**1** and tripeptides D-**4** and *R*-**11** over multiple concentrations. The Δ*frha-pbd* mutant was included as a negative control. Bacteria were incubated statically with and without inhibitors in polystyrene tubes containing lysogeny broth (LB) medium. After removing medium and planktonic cells by decanting, surface-attached biofilms were dried and stained with crystal violet (Figure 6). Biofilms were solubilized in DMSO and levels were quantified by measuring the absorbance of crystal violet (l = 570 nm).

**Figure 6.**
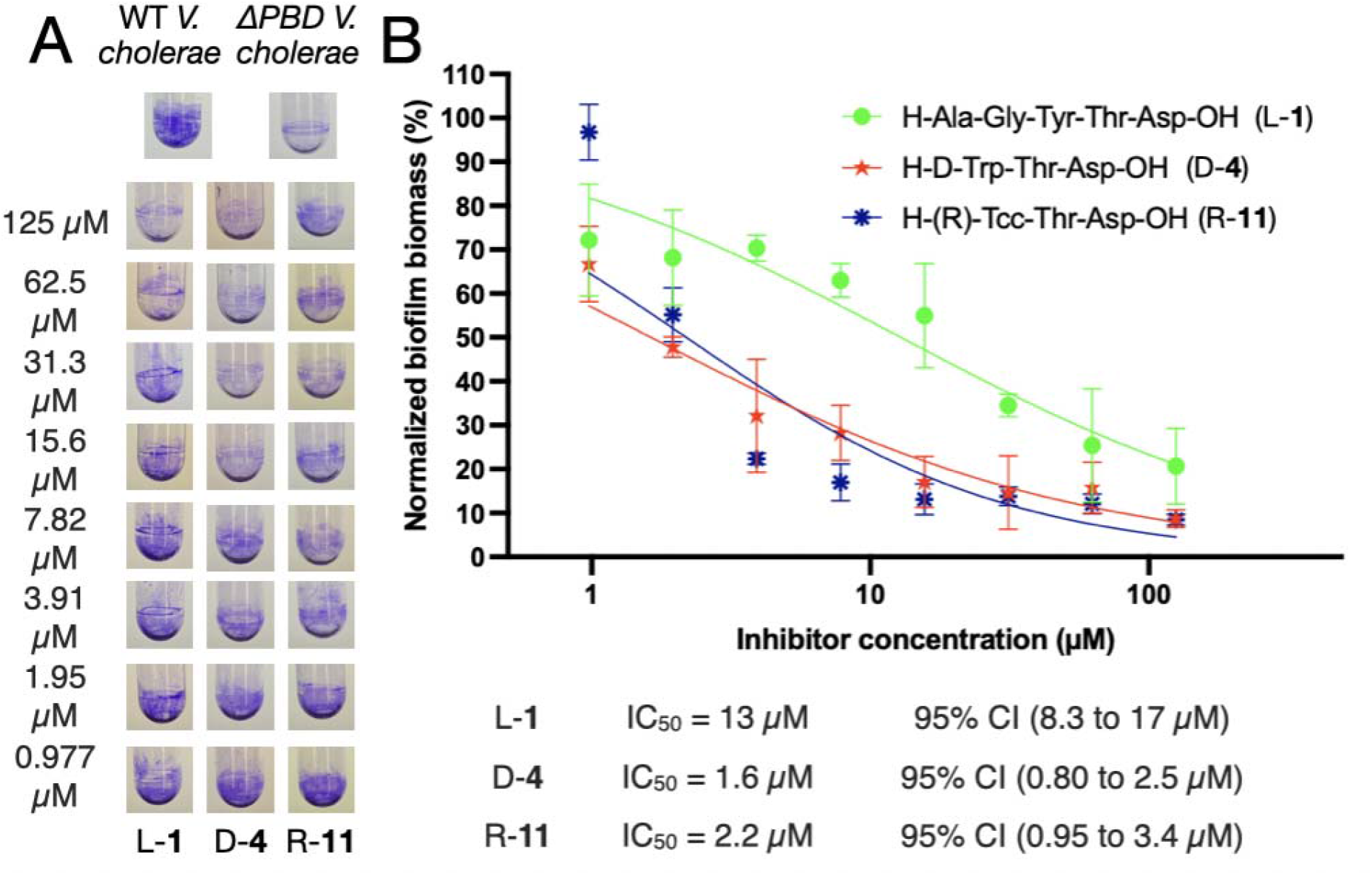
Inhibition of *Vibrio cholerae* biofilm formation. (Left) Static biofilm formation after 48 h of WT *V. cholerae* O395 and Δ*frha-pbd* deletion mutant in the presence and absence of inhibitors at indicated concentrations. (Right) After crystal violet staining, biofilms were solubilized in DMSO and levels quantified measuring the absorbance at 570 nm; ICoo values were normalized to those of WT *V. cholerae* O395 and Δ*FrhA-PBD* and ascertained using an inhibitor concentration-versus-normalized response model. Conditions were assayed in triplicate.

The biofilm was significantly reduced for the Δ*frha-pbd* mutant compared with WT *V. cholerae* O395 (A570 = 0.21 vs 0.86), consistent with the established role of the FrhA-PBD in promoting biofilm formation (Figure 5 and Table S4). Pentapeptide L-**1** inhibited biofilm formation in a concentration-dependent manner from 0.977 to 125 µM, with an IC_50_ of 13 µM (95% CI, 8.3–17 µM). Compared with pentapeptide L-**1**, the D-Trp and (R)-Tcc analogs D-**4** and R-**11** were significantly more potent, with ICoo values of 1.6 µM (95% CI, 0.80–2.5 µM) and 2.2 µM (95% CI, 0.95–3.4 µM), respectively (D-**4** vs L-**1**: F(1, 44) = 51.45, P < 0.0001; R-**11** vs L-**1**: F(1, 44) = 26.26, P < 0.0001).

### Inhibition of FrhA_Split-PBD_ Binding to Intestinal Cells

Inhibitory ability against FrhA-PBD adhesion to human intestinal epithelial Caco-2 cells was assessed as a pathologically relevant model^60–62^. Immobilized purified FrhA_Split-PBD_ was exposed to Caco-2 cells with and without inhibitors in microtiter wells (Figure 7, Figure S3, and Table S5). After incubation, nonadherent cells were removed by washing and retained cells were fixed with glutaraldehyde and stained with crystal violet. Stained cells were visualized by light microscopy. Retained dye was solubilized in acetic acid and absorbance was measured at 570 nm.

**Figure 7.**
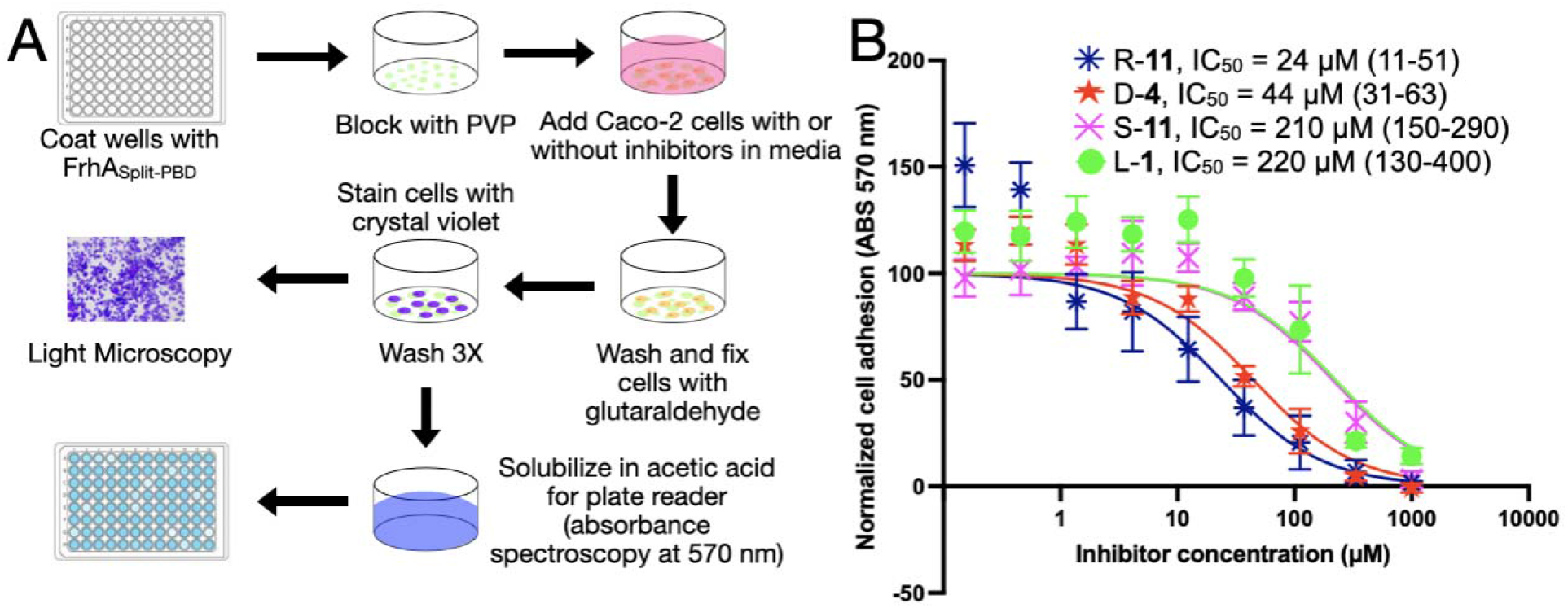
Concentration-dependent inhibition of Caco-2 cell adhesion to immobilized FrhA_Split-PBD_. (Left) Schematic of the cell adhesion assay which assesses crystal violet staining using light microscopy and quantification of retained dye measuring absorbance at 570 nm. (Right) Concentration–response curves for peptides L-**1**, D-**4**, *R*-**11**, and *S*-**11**. Symbols represent the mean ± SD of three replicate wells per concentration. Curves represent nonlinear regression fits of inhibitor concentration versus normalized response. ICoo values (95% confidence intervals): L-**1** (220 µM, 130–400), D-**4** (44 µM, 31–63), *R*-**11** (24 µM, 11–51), and *S*-**11** (210 µM, 150–290).

Concentration-dependent inhibition was evaluated for peptides L-**1**, D-**4**, *R*-**11**, and *S*-**11** over nine concentrations ranging from 0.1524 to 1000 µM. All four peptides decreased levels of adherent cells with increasing concentration (Figure 7) as observed by microscopy (Figure S3). Nonlinear regression of inhibitor concentration versus normalized adhesion yielded ICoo estimates with 95% profile-likelihood confidence intervals: L-**1**, (220 µM, 130–400 µM), D-**4**, (44 µM, 31–63 µM), *R*-**11** (24 µM, 11–51 µM), *S*-**11** (210 µM, 150–290 µM), extra sum-of-squares F-test, F_1,52_ = 30.11, P < 0.0001, Table S4. The concentration–response measurements indicated that compared to pentapeptide L-**1** and tripeptides possessing *S*-aromatic residues the *R*-tripeptide counterparts D-**4** and *R*-**11** were significantly more potent.

### Inhibitor Resistance to Chymotrypsin Proteolysis

The better potency of the peptidomimetic *R*-**11** relative to the pentapeptide L-**1** in the intestinal epithelial cell-binding assay raised the possibility that differences in proteolytic stability influenced activity, because peptidases and proteases expressed by the Caco-2 cells may degrade peptide inhibitors ^63–65^. The effects of configuration, chain length, and the Tcc residue on enzymatic proteolysis were assessed using bovine pancreatic α-chymotrypsin (100 µM) as a structural and functional homologue of human chymotrypsin B1. Chymotrypsin acts as a major pancreatic digestive protease in the small intestine, preferentially cleaving the *C*-terminal peptide bond of aromatic residues^66–70^. The relative stabilities of peptides L-**1**, L-**2**, D-**2**, L-**3**, D-**3**, L-**4**, D-**4**, *S*-**11**, and *R*-**11** towards chymotrypsin digestion were monitored using LC-MS analysis over a 120-min time course (Figures 8 and S4).

**Figure 8.**
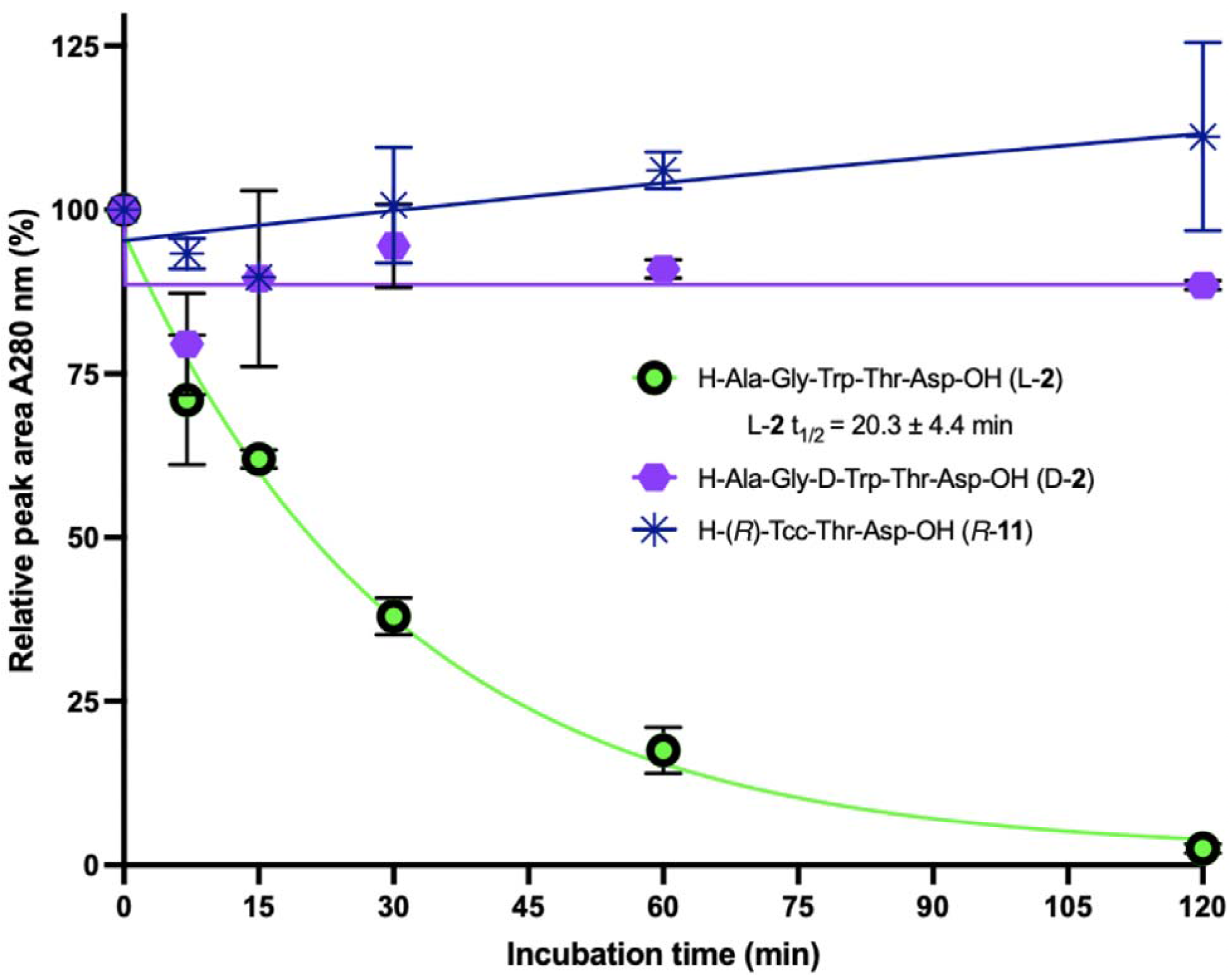
Influence of peptide structure on α-chymotrypsin proteolysis. LC–MS analysis of bovine pancreatic α-chymotrypsin digestion of inhibitors D-**2**, L-**2**, and *R*-**11** over 120 min. Peak areas measured by UV absorbance at 280 nm were normalized to the corresponding undigested controls. Each time point represents two independent experiments for D-**2** and L-**2** and three for *R*-**11**. The disappearance of L-**2** was fit with a one-phase exponential decay model to obtain a half-life of 20.3 ± 4.4 min (Tables S7 and S8).

Pentapeptides L-**1** and L-**2** with natural sequences underwent respectively progressive proteolysis to the H-Ala-Gly-Tyr-OH and H-Ala-Gly-Trp-OH cleavage products with apparent half-lives of 58.1 and 20.3 ± 4.4 min based on a one-phase exponential model (Figures 8 and S4, Tables S6 and S7). In contrast, pentapeptide D-**2** with D-tryptophan, D- and L-Tyr and Trp tripeptides D- and L-**3** and **10** as well as Tcc counterpart *R*-**11**, all exhibited negligible digestion over two hours regardless of configuration (Figures 8 and S4, Table S7). Truncation, stereochemical inversion, and the Tcc residue, all effectively conferred proteolytic stability to chymotrypsin.

## DISCUSSION

The disease cholera continues to threaten public health partly owing to emerging antibiotic resistance^12–14^. The causative pathogen *Vibrio cholerae* employs the adhesion protein FrhA to colonize the small intestine ^29^. Previously pentapeptide L-**1** was shown to bind the FrhA peptide binding domain (PBD) and competitively block bacterial adhesion in an intestinal infection model at high millimolar concentrations^29^. In a structurally guided peptidomimetic approach to develop anti-adhesion therapy based on pentapeptide L-**1**, efforts have focused on improving binding affinity, biological activity, and resistance to proteolysis in the gastrointestinal tract. Towards this goal, the co-crystal structure of H-(*S*)-Tcc-Thr-Asp-OH (*S*-**11**) bound to FrhA_Split-PBD_ is now reported providing the first experimental evidence of a tripeptide interacting within the binding pocket and further support of the importance of conserved hydrogen bonds and coordination of the two Ca^2^□ ions by the *C*-terminal Thr-Asp motif. Consistently, stereochemical and structural modifications of the L-threonine residue caused significant loss of binding affinity. On the other hand, inversion of the stereochemistry and constraint of the neighboring aromatic residue gave a set of tripeptides exhibiting increased binding affinity. Moreover, improved binding affinity translated into improved activity in functional assays. For example, D-Trp and (*R*)-Tcc peptides D-**4** and *R*-**11** exhibited respectively >2 and >3 fold improvements in inhibiting hemagglutination by *V. cholerae* than pentapeptide L-**1**^29^. Moreover, peptides D**-4** and *R***-11** blocked biofilm at low (1.6-2.2 mM) concentrations, 6 to 8-fold better than pentapeptide L-**1**. In a novel intestinal cell-binding assay, peptides D**-4** and *R***-11** were blocked Caco-2 cell adhesion to FrhA-PBD at low (24 and 44 mM) concentrations 5 to 9-fold better than pentapeptide L-**1**. Notably, nanomolar binding affinities measured with purified proteins translated into micromolar inhibition of hemagglutination, biofilm formation, and Caco-2 cell adhesion. Considering factors responsible for this disparity, proteolytic stability was examined in a chymotrypsin stability assay in which natural pentapeptides L-**1** and L-**2** were readily cleaved *C*-terminal to the respective L-Tyr and L-Trp residues, but the corresponding D-Trp pentapeptide D-**2** proved stable. Moreover, abridgement to the tripeptide and application of the Tcc residue significantly impeded chymotrypsin proteolysis.

The binding affinity and activity of the described ligands across hemagglutination, biofilm, and intestinal cell adhesion assays support further development of FrhA-PBD antagonism as an anti-adhesion strategy against *V. cholerae*. On the path toward an effective drug candidate, further development is required, including evaluation of cytotoxicity, appropriate dosage, and local exposure. Our findings provide a foundation and justification for studies in animal models. Continued development of antagonists of FrhA, alone and in combination with other virulence factor inhibitors, offers promise for disrupting the pathogenesis caused by pandemic *V. cholerae*.

## EXPERIMENTAL SECTION

### Protein expression and purification for X-ray crystallography and assays

The FrhA_Split-PBD_ construct was expressed according to previously published protocols^41, 71, 72^ in *Escherichia coli* BL21(DE3) (Thermo Fisher) cells carrying an expression vector based on pET28a. The protein was purified based on previously published protocols^41^ with a Ni-NTA affinity chromatography column and a HiLoad™ 16/60 Superdex 200 pg (Cytiva) size-exclusion column.

### Peptide synthesis

The compounds L-**1**, L-**2**, D-**2**, L-**3**, D-**3**, L-**4**, D-**4**, **5**, **6**, **7**, **8**, **9**, and **10** were purchased from GenicBio (Shanghai, China) as lyophilized peptide powders. LC-MS performed by the vendor supported a purity of >95% (Figure S5).

### Synthesis of Tcc analogues

#### General

Unless otherwise specified, reagents from commercial sources were used as received. Dichloromethane (DCM), dimethylformamide (DMF), water (H_2_O), and methanol (CH_3_OH) were obtained from Fisher Chemical. Amino acids, coupling reagents, and CTC resin were obtained from Chem-Impex, Combi-Blocks, Oakwood Chemicals, and Ambeed.

Liquid Chromatography-Mass Spectrometry (LC-MS) was performed using an Agilent series symmetry C18 column (3.5 μm, 4.6 × 75 mm), maintained between 27 - 30 °C, and 118.7 - 141.5 bar, using a flow rate of 0.8 mL min^-1^, MSD parameters, an ES-API ionization source, positive ionization polarity, and 5.0 V ionization energy. Mass spectrometric data are reported in the form of m/z (intensity relative to the base peak = 100). The UV detection employed a diode array detector. The mobile phase consisted of solvent I, H□O containing 0.1% formic acid (FA) and solvent II, CH_3_CN containing 0.1% FA or solvent III, CH_3_OH containing 0.1% FA. The elution gradients over 14 min were as follows: Method A: 5–50% II in I; Method B: 5–50% III in I.

#### Representative protocols for the activation and loading of CTC resin and the determination of the resin loading

##### Preparation of Fmoc-Asp(O*t*-Bu)-O-2-CTC

CTC resin (1.90 g, 0.3-0.8 mmol/g, 75-100 mesh) was swollen for 10-15 min in 20 mL of DCM in a polystyrene tube fitted with Teflon™ filter, stopper, and stopcock. The resin mixture was filtered. The resin was treated with a 1:4 SOCl_2_: DCM solution (20 mL), shaken vigorously for 60 min on a RotoMix50800 automated shaker, filtered, and washed with DCM (5 x 20 mL for 2-3 min/wash). The resin was treated with a solution of Fmoc-Asp(O*t*-Bu)-OH (1.19 g, 2.89 mmol, 200 mol%) in DCM (20 mL) followed by DIPEA (629 μL, 3.61 mmol, 250 mol%), agitated for 60 min on the automated shaker, filtered, capped by agitation for 15 min in HPLC-grade methanol (2 mL), filtered, washed sequentially with DCM, DMF, and MeOH (3 x 5 mL for 2-3 min/wash), and dried in vacuo for three h. Loading measurement was done according to a reported procedure^57^; a 10 mg aliquot of resin was swollen in DMF (0.2 mL), treated twice with a 20% piperidine in DMF solution (0.2 mL), and filtered. The filtrates were combined, transferred into a 25 mL volumetric flask, and diluted to full volume with ethanol. The ethanol solution of the Fmoc-cleaved resin was compared with blank (0.4 mL of 20% piperidine in DMF diluted similarly with EtOH) and standard (3 mg, of Fmoc-Asp(O*t*-Bu)-OH treated with 0.4 mL of 20% piperidine in DMF diluted similarly with EtOH), and the loading was calculated to be 0.4 mmol/g based on comparisons of the UV measurement at 301 nm.

#### Amino acid couplings, Fmoc deprotections and CTC resin cleavage

Peptide synthesis was performed on CTC resin in plastic filtration tubes equipped with a polyethylene filter and stopcock with agitation on an automated shaker. After Fmoc group removal from the peptide by treatment with 20% piperidine in DMF (2 mL/100 mg of resin) for 30 min, the resin gave a positive test^73^. Couplings were performed by premixing the protected amino acid (300 mol%) and HBTU (300 mol%) as the coupling agent in DMF (2 mL/100 mg of resin) for five minutes, treating with Hünig’s base (DIPEA, 600 mol%), and transferring with a syringe to the resin. The resin mixture was agitated on a mechanical shaker. The completion of the coupling reaction was confirmed by a negative Kaiser test and by LC-MS analysis of a resin aliquot cleaved with 20% HFIP in DCM. After each coupling and deprotection step, the resin was washed using a sequence of 3-4 min per wash and two mL of the following solvents per 100 mg of resin: DMF (2×), MeOH (2×), and DCM (2×). The resin was cleaved for 1 h and washed using 20% HFIP in DCM followed by DCM. The filtrates and washing were combined and evaporated under reduced pressure.

#### H-(*S*)-Tcc-Thr-Asp-OH (*S*-11)

Fmoc-Asp(O*t*-Bu)-O-2-CTC (loading. = 0.4 mmol/g, 1.90 g, 1.44 mmol, 100 mol %) was swollen with 20% piperidine in DMF and shaken for 30 min to remove the Fmoc protecting group. After filtration, washing, and drying, the resin was swollen in DMF (5 mL), treated subsequently with a pre-mixed solution of Fmoc-Thr(O*t*-Bu)-OH (1.72 g, 2.28 mmol, 300 mol%), HBTU (865 mg, 2.28 mmol, 300 mol%), and DIPEA (0.794 mL, 4.56 mmol, 600 mol%) in DMF (20 mL), and agitated for 3h. Cleavage of a resin aliquot with 20M% HFIP in DCM and LC-MS analysis of the cleaved product at 254 nm confirmed complete conversion to the dipeptide Fmoc-Thr(O*t*-Bu)-Asp(O*t*-Bu)-OH. The resin was filtered and washed. Fmoc deprotection was carried out by shaking the resin with 20% piperidine in DMF for 30 min. After filtration, washing, and drying, the resin was swollen in DMF (20 mL) and treated with a pre-mixed solution of Fmoc-Trp-OH (972 mg, 2.28 mmol, 300 mol%), HBTU (865 mg, 2.28 mmol, 300 mol%), and DIPEA (0.794 mL, 4.56 mmol, 600 mol%) in DMF (20 mL). After cleavage of a resin aliquot with 20M% HFIP in DCM, LC-MS analysis at 254 nm confirmed complete conversion to the tripeptide Fmoc-Trp-Thr(O*t*-Bu)-Asp(O*t*-Bu)-OH. The resin was filtered and washed. Fmoc deprotection was carried out on 100 mg of the tripeptide using a 20% piperidine in DMF and shaken for 30 min.

After filtration, washing, and drying, H-Trp-Thr(O*t*-Bu)-Asp(O*t*-Bu)-O-CTC (**12**, 100 mg, 0.38 mmol) was swollen in DMF for five minutes, treated with CH_2_O (37% aq. solution, 5.66 μL, 0.38 mmol, 100 mol%) and PPTS (28.4 mg, 0.19 mmol, 50 mol%), and agitated for 18 h. A resin aliquot was cleaved and analyzed by LC-MS at 254 nm to confirm completion of cyclization. The resin was dried and cleaved using 20% HFIP in DCM to give H-(*S*)-Tcc-Thr(O*t*-Bu)-Asp(O*t*-Bu)-OH **14**, which was treated with 50% TFA in DCM (3 mL) for 1h. The volatiles were evaporated. The residue was purified by reversed-phase HPLC on a Waters C18 column (SunFire™, particle size, 10 μm, 150 mm × 10 mm) using a linear gradient of 30–45% CH_3_CN (0.1% FA) in water (0.1% FA) with a flow rate of 4.2 mL/min over 12 min. Freeze drying of the collected fractions gave H-(*S*)-Tcc-Thr-Asp-OH (S-**11**) as a brown solid (6.3 mg, 21 %). RP-HPLC, RT = 3.362 min [Method C, >95% purity]; RT = 4.810 min [Method D, >95% purity].

#### H-(*R*)-Tcc-Thr-Asp-OH (*R*-11)

Employing the procedure described for *S*-**11,** H-D-Trp-Thr(O*t*-Bu)-Asp(O*t*-Bu)-O-CTC (**12**, 100 mg, 0.38 mmol) was swollen in DMF for five minutes, treated with CH_2_O (37% aq. solution, 5.66 μL, 0.38 mmol, 100 mol%) and PPTS (28.4 mg, 0.19 mmol, 50 mol%), and agitated for 18 h. A resin aliquot was cleaved and analyzed by LC-MS at 254 nm to confirm completion of cyclization. The resin was dried and cleaved using 20% HFIP in DCM to give H-(*R*)-Tcc-Thr(O*t*-Bu)-Asp(O*t*-Bu)-O-CTC **14** which was treated with 50% TFA in DCM (3 mL) for 1h. The volatiles were evaporated. The residue was purified by reversed-phase HPLC on a Waters C18 column (SunFire™, particle size, 10 μm, 150 mm × 10 mm) using a linear gradient of 30–45% CH_3_CN (0.1% FA) in water (0.1% FA) with a flow rate of 4.2 mL/min over 12 min. Freeze drying of the collected fractions gave H-(*R*)-Tcc-Thr-Asp-OH (*R*-**11**) as a brown solid (6.3 mg, 19%). RP-HPLC, RT = 3.362 min [Method C, >95% purity]; RT = 4.810 min [Method D, >95% purity].

#### Microscale thermophoresis

MST was performed in the presence of the fluorescently labeled FrhA_Split-PBD_ using unlabeled ligands based on a previously published protocol^41^ with the following modifications. The concentration of fluorescently labeled FrhA_Split-PBD_ ranged from 80 nM to 20 nM (Table S8) to accommodate the high binding affinity of ligands. A series of sixteen 1:1 dilution of the ligands was prepared for each ligand to place the expected *K*_d_ values in the logarithmic center of the concentration data points (Table S8). Experiments were performed in triplicate, with the K_d_ reported as the mean and its error as the standard deviation of the three independent experiments. In the NanoTemper MO.Affinity Analysis software (version 2.3, NanoTemper Technologies), the *K*_d_ values were determined by plotting the change in normalized fluorescence [ΔF_norm_(‰)□=□F_1_/F_0_] against the logarithm of the concentrations of peptide ligands, where F_1_ and F_0_ correspond to the heated and baseline regions of the thermophoresis traces, respectively^74^. The time between F and F_0_ was set to 20□s for analysis.

#### X-ray crystallography and model building

FrhA_Split-PBD_ was crystallized with (S)-Tcc-Thr-Asp by the microbatch method. Each 5.0 µL crystallization drop contained 2.0 µL of crystallization solution (0.2 M calcium acetate, 0.1 M HEPES, pH 7.5, and 10% (w/v) PEG 8000), 2.0 µL of FrhA_Split-PBD_ (9.7 mg/mL in 50 mM Tris-HCl, pH 9.0, 200 mM NaCl, and 5 mM CaCl□), 0.5 µL of 4.5 mM (S)-Tcc-Thr-Asp, and 0.5 µL of 5% polyvinylpyrrolidone K15 (PVP K15). Thick, hexagonal crystals formed after approximately three weeks. Diffraction data were collected at the CMCF-ID beamline of the Canadian Light Source synchrotron. The phase problem was solved using molecular replacement with the structure of this construct in complex with L-**1** (PDB ID: 9YBQ). The initial models were improved by rounds of refinement in Phenix ^75, 76^ and manual model building in Coot ^77, 78^. The atomic model and restraints for the ligand S-**11** were generated using AceDRG in the CCP4 suite and were used for model building and refinement of the final X-ray crystal structure^79^. Figures relevant to structural studies were prepared using ChimeraX 1.10 ^80, 81^. For details, see Table S2.

#### Bacterial hemagglutination and inhibition assay

This assay was adapted from previously published protocols^28, 29, 82^. WT and Δ*FrhA-PBD V. cholerae* O1 O395 strains were grown overnight in LB medium supplemented with 100 µg/mL streptomycin. Inhibitors were serially diluted 1:1 seven times to give eight concentrations ranging from 4.0 mM to 31 µM in a round-bottomed 96-well microtiter plate, with each well holding 50 µL of inhibitor solutions. Bacteria were washed and resuspended to an OD600 of 0.5 in KRT buffer (128 mM NaCl, 5.1 mM KCl, 1.3 mM MgSO□, 7.7 mM Tris-HCl (pH 7.4), and 2.75 mM CaCl□)^82^. Aliquots of 100 µL of bacterial suspension were mixed with aliquots of 50 µL inhibitor solutions. Human type O red blood cells (RBCs; sourced from the McGill University Health Centre) were washed three times and resuspended in KRT buffer. An aliquot (50 μL) of 2% red blood cell suspension in KRT buffer was added to 150 μL of bacterial-inhibitor suspension. In RBC-only and inhibitor-free control groups, 150 µL or 50 µL KRT buffer was added, respectively. Plates were incubated statically at room temperature and monitored for HA titer at two h. The grayscale image of the plate was analyzed by ImageJ to determine the IC_50_ according to a published protocol^48^. The HA experiments were performed a minimum of three separate times, and results from all three experiments are shown in the Supplemental Information.

#### Crystal violet *V. cholerae* biofilm assay

This assay was adapted from previously described assays^28, 29, 83, 84^. Overnight cultures of *V. cholerae* (WT and ΔFrhA-PBD *V. cholerae* O1 O395) in LB medium with 100 µg/ml added streptomycin were centrifuged and resuspended in fresh LB medium with streptomycin. Sterile, capped 10 mL polystyrene culture tubes were filled with 0.75 mL of LB broth containing *V. cholerae*. The bacterial suspension was adjusted to a final OD600 of 0.10, and inhibitors in LB medium were added at the indicated final concentrations. Samples were inoculated in triplicate and allowed to incubate statically at 37 °C for 48 h.

After incubation, the tubes were air-dried for 12 h. Adherent bacteria and biofilm matrix were stained with crystal violet and rinsed three times with Milli-Q water. To solubilize attached crystal violet, 1 mL of dimethyl sulfoxide (DMSO) was added to each tube. Biofilm formation was quantified by transferring 100 µL of each DMSO-solubilized sample to a microtiter plate and measuring absorbance at 570 nm.

#### Crystal violet Caco-2 cell-binding assay

This assay was adapted from previously described assays^60, 85^. To test the binding of the FrhA_Split-PBD_ construct to human intestinal epithelial cells and to quantify the potency of the inhibitors, 96-well plates (Maxisorp, Nunc) were coated with protein. The wells were coated with 100□μl of either FrhA_Split-PBD_ construct (100□μg/ml) or 1% (w/v) polyvinylpyrrolidone (Sigma) in TBS-Ca^2+^ buffer (50 mM Tris-HCl pH 9, 200 mM NaCl, 5 mM CaCl_2_) overnight at 4°C. Non-specific binding sites were blocked with 100□μl 1% (w/v) polyvinylpyrrolidone 10 (PVP) in TBS-Ca^2+^ buffer for 1□h at 20°C. Caco-2 cells grown to 90% confluency in DMEM or RPMI 1640 medium (Gibco) were briefly treated with trypsin to detach them from the flask, after which the trypsin was inactivated with 1% serum in RPMI 1640 medium. About 20,000 Caco-2 cells, premixed with inhibitors (when applicable) solubilized in RPMI 1640 medium to the assay concentrations, were added to each well and incubated for 1 h in RPMI 1640 medium at 37°C. Non-attached cells were removed by washing with 200 µL of TBS-Ca^2+^ buffer with three gentle taps and inverting the plate. Attached cells were fixed with 5% (v/v) glutaraldehyde in TBS-Ca^2+^ buffer and stained with 0.1% (w/v) crystal violet (Sigma) in MES buffer (200 mM 2-morpholin-4-ylethanesulfonic acid, pH 6, Sigma) for 30□min. After the wells were washed three times with H_2_O, the level of cell adhesion was also assessed by light microscopy (Nikon Eclipse Ts2, Nikon, Japan and B-290TB, Optika, Italy). The dye was solubilized in 100 µL 10% (v/v) acetic acid (BioShop, Canada) and quantified at 570□nm using a plate reader (BioTek Synergy H1M2, BioTek, USA) for quantification of Caco-2 cell adhesion. Each condition used three wells. FrhA_Split-PBD_-coated wells without inhibitor served as the positive control, whereas wells treated with PVP 10 alone served as the negative control. Responses were normalized to the arithmetic mean signals of these controls, defined as 100% and 0% adhesion, respectively.

#### Chymotrypsin digestion assay

Peptides were prepared as 2 mM working solutions in Milli-Q water. For each digestion reaction, 125 µL of peptide solution was transferred to a microcentrifuge tube. Lyophilized bovine pancreatic α-chymotrypsin (Sigma-Aldrich, C4129) was freshly dissolved in TBS-Ca^2^□ buffer (50 mM Tris-HCl, pH 7.5, 150 mM NaCl, and 5 mM CaCl□) to a concentration of 200 µM and kept on ice until addition to incubation tubes. Proteolysis was initiated by mixing 125 µL of the peptide solution with 125 µL of the α-chymotrypsin solution, yielding final concentrations of 1 mM peptide and 100 µM α-chymotrypsin in a 250 µL reaction volume. Samples were incubated at 37 °C for 7, 15, 30, 60, or 120 min in a Fisher Scientific Isotemp 500 incubator. Chymotrypsin-free control samples with an equal substrate concentration were prepared by adding a volume of 125 µL Milli-Q water in place of α-chymotrypsin. Proteolysis was quenched by the addition of 13.5 µL of 98% formic acid (FA; Fluka Analytical) to each sample, yielding a final volume of 263.5 µL and a formic acid concentration of approximately 5% (v/v). Samples were mixed by vortexing, flash-frozen in liquid nitrogen, and stored at −80 °C until liquid chromatography–mass spectrometry (LC–MS) analysis.

Post-proteolysis samples were analyzed by LC–MS using an Agilent Technologies system with an Agilent C18 column (3.5 µm, 4.6 × 75 mm; Agilent Technologies). The aqueous mobile phase consisted of water containing 0.1% FA, and the organic mobile phase consisted of methanol containing 0.1% FA. Chromatographic separation was performed using a 5–50% MeOH (+ 0.1% FA) in H_2_O (+ 0.1% FA) gradient over 12 min at a flow rate of 0.8 mL/min. Mass spectrometric detection was performed using an electrospray ionization source operated in positive-ion mode. Proteolysis was assessed by monitoring ions corresponding to the intact peptide and its cleavage products. Extracted-ion chromatograms were generated using Agilent software, and the corresponding chromatographic peaks were individually integrated. Integrated signals were used to assess changes in the abundance of intact substrate and cleavage products over the digestion time course. For quantification of intact peptide remaining, chromatographic peaks containing protonated molecular ions corresponding to the intact peptide were summed. The percentage of intact peptide remaining was plotted as a function of time and fitted to a one-phase exponential decay model using GraphPad Prism, with the signal from the chymotrypsin-free control normalized to 100%. Peptide half-life (t□/□) was calculated from the fitted decay curve. Each time point comprised measurements from two independent experiments.

## ACKNOWLEDGMENTS

We thank Peter L. Davies from Queen’s University for fruitful discussions. We are grateful to Mindy Kang, Ahoor Saleem, Ziran Gan, Sasha Tan, and Justine Dela Cruz for their assistance with protein purification and other technical assistance. We thank staff at the CMCF-ID beamline at the Canadian Light Source (CLS) for access and assistance with remote X-ray crystal diffraction data collection. We thank Anirudh Mantri and Martin Schmeing at McGill University for sharing access to collecting X-ray crystallography data at the CLS CMCF-ID beamline. We are grateful for a doctoral research scholarship from the Fonds de recherche du Québec (FRQ; grant 2006473; https://doi.org/10.69777/2006473) awarded to M.W. We are grateful for funding from the Natural Sciences and Engineering Research Council of Canada (NSERC, Discovery Research Project RGPIN-2025-04423), the Canadian Institutes of Health Research (CIHR, Project PJT-186296), and the Fonds de Recherche du Québec - Nature et Technologie for support of the Biodiversa+ program (project # 356910) and for funding the Centre in Green Chemistry and Catalysis (CGCC, FRQNT-2020-RS4-265155-CCVC). Assistance is acknowledged from members of the Université de Montréal facilities: Dr. P. Aguiar (NMR spectroscopy) and Dr. A. Fürtös (mass spectrometry). Operation of the NMR spectrometers at the Regional Centre for Magnetic Resonance in the Department of Chemistry at the Université de Montréal was made possible through funding from the Canada Foundation for Innovation and the Institute Courtois.

## Author Information

### Author Contributions

The manuscript was written through contributions of all authors. All authors have given approval to the final version of the manuscript.

### Notes

The authors declare no competing financial interests.

## ASSOCIATED CONTENT

The Supporting Information is available free of charge.

Accession Codes: The PDB code for *S*-**11** is 37YK. The authors will release the atomic coordinates and experimental data upon article publication.

## ABBREVIATIONS

Alt: allothronine
ANOVA: analysis of variance
CCP4: Collaborative Computational Project Number 4 in Protein Crystallography
CI: confidence interval
CMCF-ID: Canadian Macromolecular Crystallography Facility – Insertion Device
CTC: 2-chlorotrityl chloride
DCM: dichloromethane
DIPEA: *N,N*-diisopropylethylamine
DMEM: Dulbecco’s Modified Eagle Medium
DMF: dimethylformamide
DMSO: dimethyl sulfoxide
ES-API: electrospray atmospheric pressure ionization
EtOH: ethyl alcohol
FA: formic acid
FrhA: Flagellar Regulated Hemagglutinin A
Fmoc: 9-fluorenylmethoxycarbonyl protecting group
HA: hemagglutination
HBTU: 2-(1H-benzotriazol-1-yl)-1,1,3,3-tetramethyluronium hexafluorophosphate
HEPES: 4-(2-hydroxyethyl)-1-piperazineethanesulfonic acid
HFIP: Hexafluoro-2-propanol
HPLC: High-performance liquid chromatography
IC□□: Half maximal inhibitory concentration
Kd: dissociation constant
Hyp: hydroxyproline
KRT: Krebs-Ringer-Tris buffer
LB: lysogeny broth
LC-MS: liquid chromatography–mass spectrometry
MeCN: Acetonitrile
MeOH: methanol
MES: 2-(morpholin-4-yl)ethanesulfonic acid
MpIBP: Marinomonas primoryensis ice-binding protein
MSD: mass selective detector
MST: microscale thermophoresis
NTA: Nitrilotriacetic acid
PBD: peptide-binding domain
PDB: protein data bank
PEG: polyethylene glycol
PPTS: pyridinium p-toluenesulfonate
PVP: polyvinylpyrrolidone
RBC: red blood cell
RP-HPLC: Reversed-phase high performance liquid chromatography
RT: retention time
SD: standard deviation
SOCl2: Thionyl chloride
TBS: Tris-buffered saline
Tcc: 1,2,3,4-tetrahydro-β-carboline-3-carboxylic acid
TFA: trifluoroacetic acid
UTI: urinary tract infection
UV: ultraviolet
WT: wild type

## Notes

### Competing Interest Statement

The authors have declared no competing interest.

### Summary of Updates

Compounds were renumbered. Associated figures, tables, and supplemental information were updated. Introduction, results, and discussion were edited.

