## Supplemental Information for "A peptidomimetic inhibitor of *Vibrio cholerae* adhesin protein FrhA"

Supplemental Information for
A peptidomimetic inhibitor blocks binding of *Vibrio cholerae* adhesin FrhA to intestinal cells.

Mingyu Wang^1,*^, Charity Yongo-Luwawa^2,*^, Laura Zhang^1,*^, Ryu Williston^1^, John Presley^1^, Zhaleh Mehboodi^3^, Dieter Reinhardt^1^, Karl E. Klose^4^, William D. Lubell^2,#^, Shuaiqi Guo^1,#^

1. Department of Anatomy and Cell Biology, McGill University

2. Département de chimie, Université de Montréal

3. McGill University Health Centre

4. Department of Molecular Microbiology and Immunology, University of Texas at San Antonio

*Equal contributions, ^#^Corresponding Authors

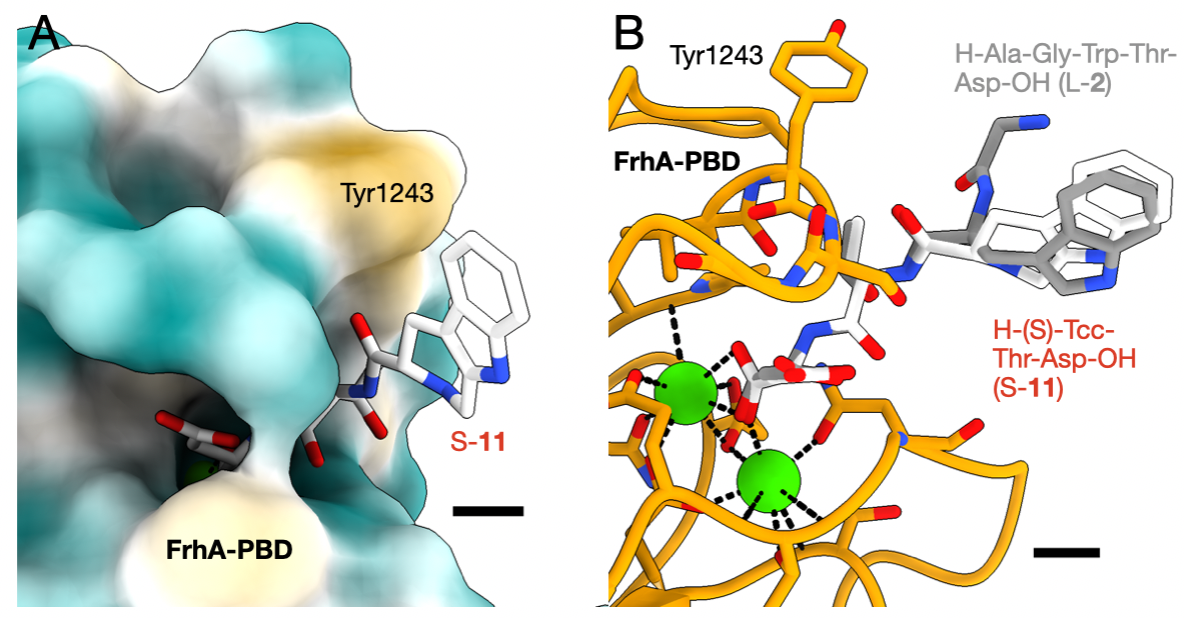

**Figure S1**: Alternative Views and LigPlot diagram of the X-ray crystal structure of S-**11** in complex with FrhA_Split-PBD_ (PDB ID: 37YK). (A) The surface of the ligand-binding pocket is shown with hydrophobic regions in yellow and hydrophilic regions in blue. (B) The binding-conformation of L-**2** (grey) superimposed on S-**11** (white) through the alignment of the FrhA_Split-PBD_ molecules in the structures 37YK and 9YBQ.

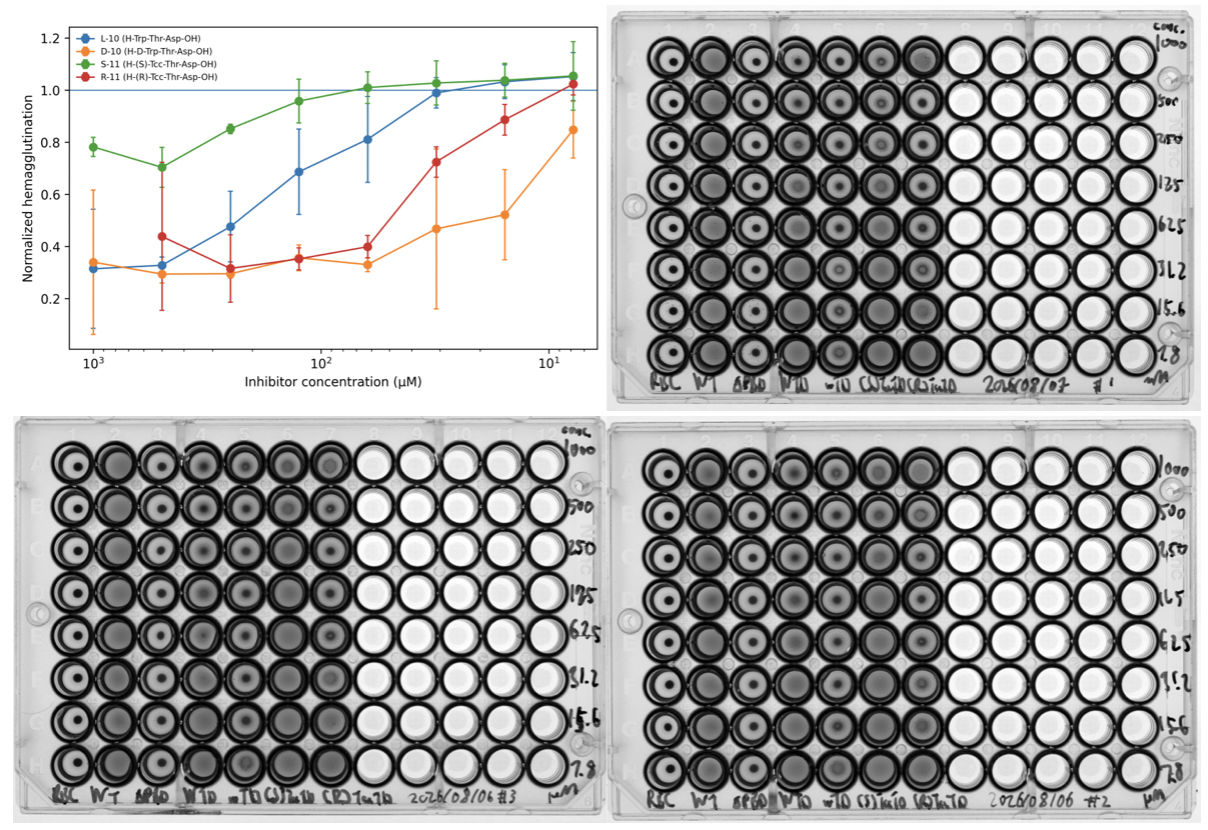

**Figure S2**: Plotted normalized hemagglutination over inhibitor concentration of the hemagglutination assay in triplicate and scans of 96-well round-bottom plates used in image analysis.

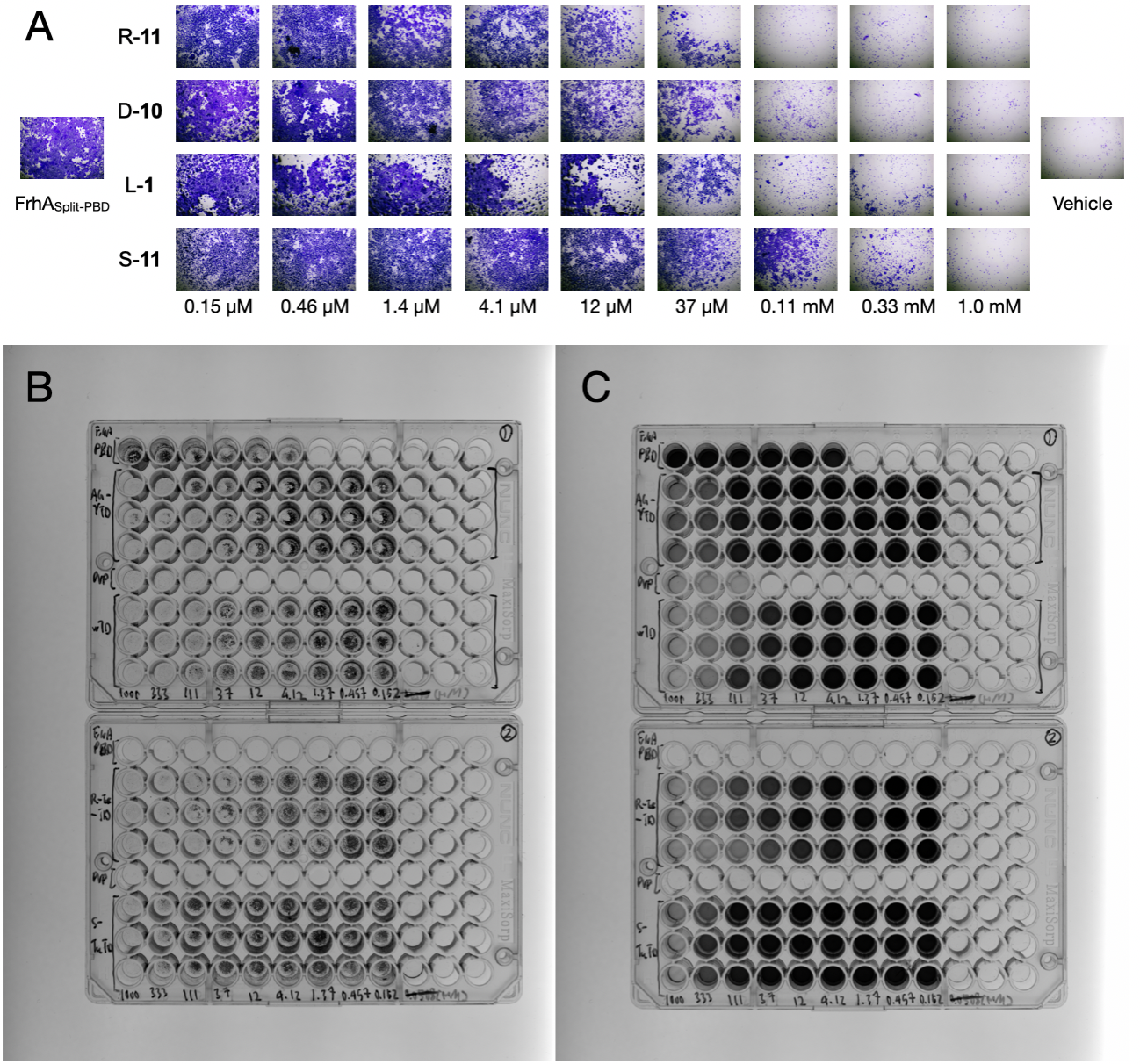

**Figure S3**: Dose-response inhibition of Caco-2 cell adhesion to immobilized FrhA_Split-PBD_ by R-**11**, D-**10**, L-**1**, and S-**11**. (A) Representative light micrographs of crystal violet-stained cells at the indicated inhibitor concentrations. (B) Image of 96-well plates after crystal violet staining and before the addition of acetic acid. (C) Images of 96-well plates after the addition of acetic acid to solubilize the retained dye. Wells A4, A5, and A6 on plate 1 contained Caco-2 cells bound to immobilized FrhA_Split-PBD_ without polyvinylpyrrolidone (PVP 10) blocking. These wells served as controls for the blocking step and were excluded from quantitative analysis.

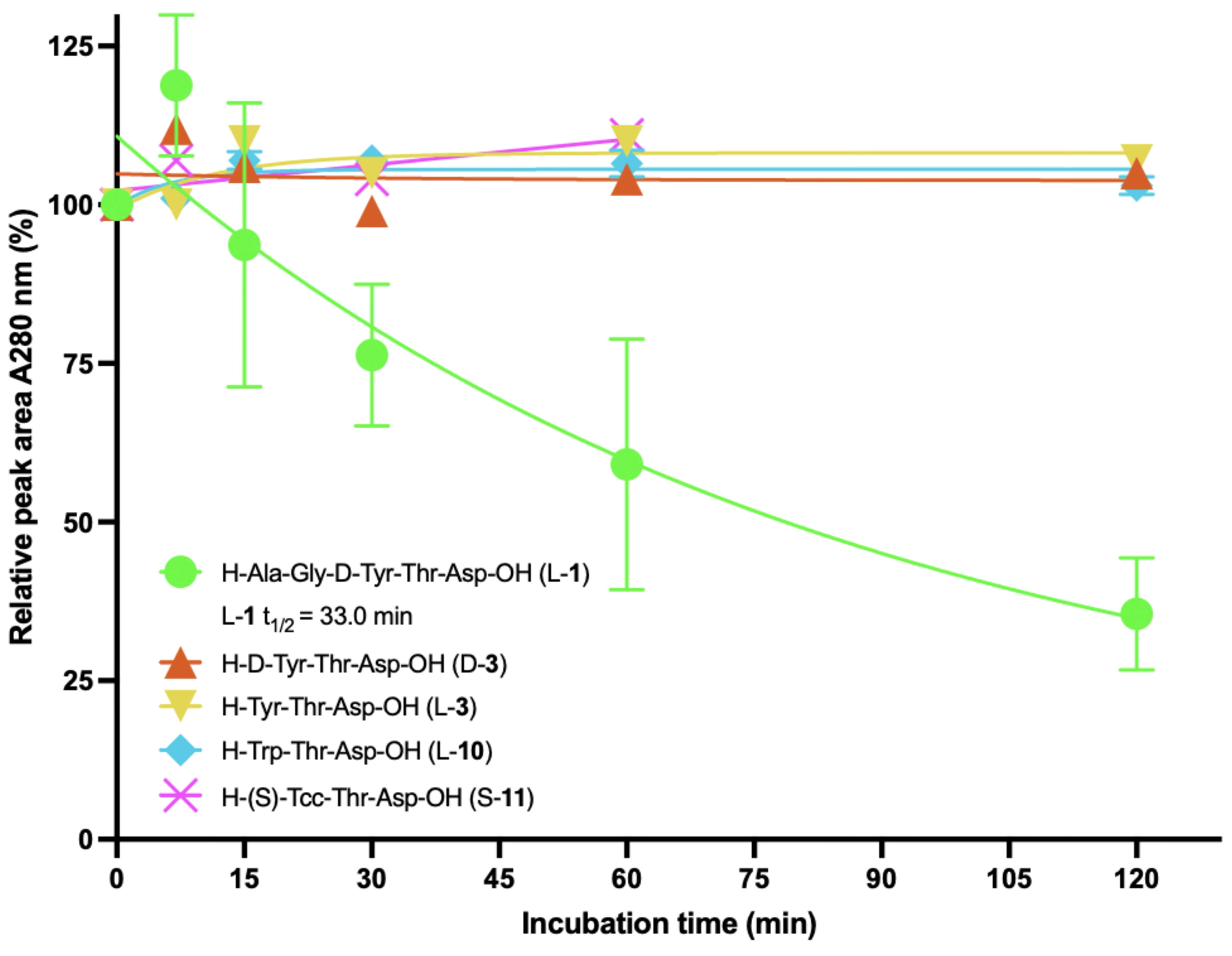

**Figure S4.** Chymotrypsin digestion of L-**1** (AGYTD) and other inhibitors. LC–MS analysis of bovine pancreatic α-chymotrypsin digestion L-**1** (green circle), D-**3** (orange triangle), L-**3** (yellow inverted triangle), L-**10** (cyan diamond) and S-**11** (pink cross) over 120 min. Peak areas measured by UV absorbance at 280 nm were normalized to the corresponding undigested controls. Each time point represents two independent experiments for L-**10** and one experiment for all others. Fitting the disappearance of L-**1** to a one-phase exponential decay model yielded a half-life of 33.0 min (Table S6).

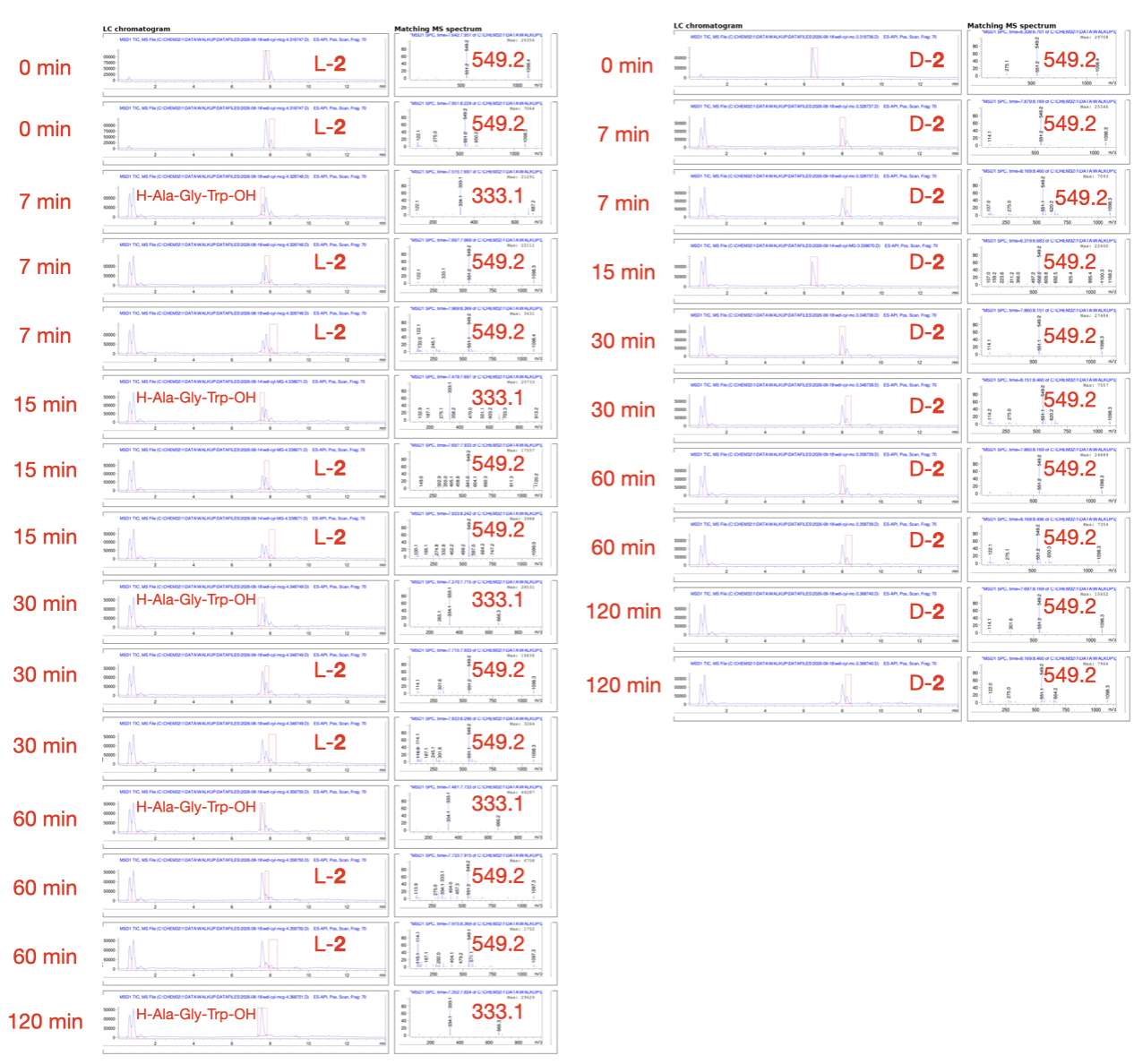

**Figure S5**: Collage of Representative chromatogram and matching mass spectra of the peak indicated by red box for the chymotrypsin digestion assay of L-**2** (left) and D-**2** (right). The digestion time is labeled on the left of each unit. The dominant species in the corresponding peak is labeled in the LC section, and the dominant m/z ratio is labeled in the MS section.

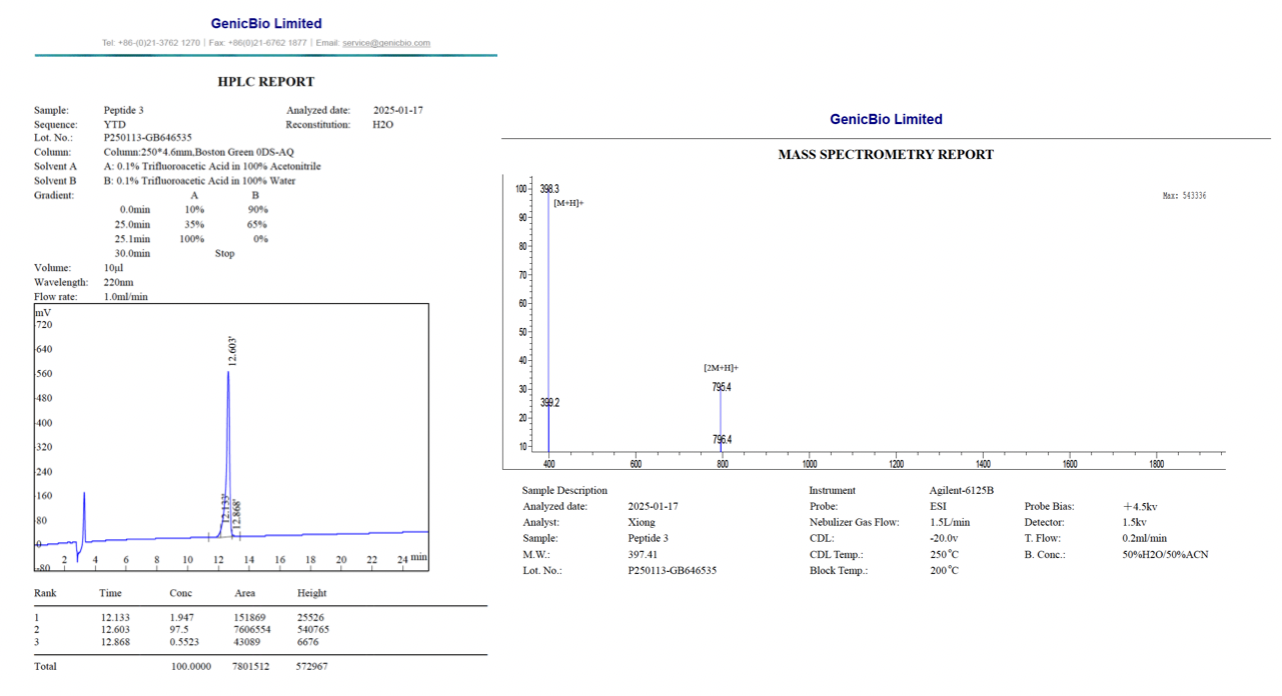

**Figure S6**: Representative chromatogram (left) and matching mass spectra (right) provided by the Vendor GenicBio Limited for compound L-**3**.

**Table S1.** Retention times, crude purity, final purity, yields and mass spectrometric data for Tcc peptides and analogues

| **Compound** | **RT (Min)** | | **Crude purity**  **%** | **Final purity**  **at 280 nm**  **%** | **Yield %** |
| --- | --- | --- | --- | --- | --- |
|  | **CH_3_OH** | **CH_3_CN** |  |  |  |
| *S*-**11** | 3.362 | 4.810 | 66 | 95 | 21 |
| *R*-**11** | 5.460 | 6.635 | 70 | 95 | 17 |

Isolated purity ascertained by LC-MS using **Method A or Method B**.

**Table S2. X-ray data collection and refinement statistics for FrhA_Split-PBD_ in complex with (S)-Tcc-Thr-Asp (PDB ID: 37YK)**

| Parameter | FrhA PBD-Split–(S)-Tcc-Thr-Asp |
| --- | --- |
| Data collection |  |
| Space group | P6₃22 |
| Unit-cell parameters |  |
| a, b, c (Å) | 119.456, 119.456, 125.008 |
| α, β, γ (°) | 90.00, 90.00, 120.00 |
| Wavelength (Å) | 0.953716 |
| Resolution range (Å) | 125.01–2.02 (2.09–2.02) |
| Total observations | 1,113,715 (38,258) |
| Unique reflections | 35,040 (3,359) |
| Completeness (%) | 99.9 (98.0) |
| Multiplicity | 31.8 (11.4) |
| ⟨I/σ(I)⟩ | 12.1 (0.9) |
| Rmerge | 0.208 (2.284) |
| Rmeas | 0.211 (2.395) |
| Rpim | 0.035 (0.691) |
| CC1/2 | 0.994 (0.317) |
| Anomalous completeness (%) | 99.7 (97.7) |
| Anomalous multiplicity | 17.0 (6.0) |
| Refinement |  |
| Resolution range (Å) | 62.50–2.20 |
| Reflections used in refinement | 27,317 |
| Working-set reflections | 25,982 |
| Rfree test-set reflections | 1,335 |
| Rfree test set (%) | 4.89 |
| Rwork | 0.200 |
| Rfree | 0.242 |
| Coordinate error, maximum-likelihood estimate (Å) | 0.22 |
| Model composition |  |
| Protein residues | 313 |
| Ligand residues, Tcc-Thr-Asp | 3 |
| Calcium ions | 6 |
| Water molecules | 303 |
| Non-H atoms, total | 2,627 |
| Protein non-H atoms | 2,287 |
| Ligand non-H atoms | 31 |
| Ca²⁺ ions | 6 |
| Water O atoms | 303 |
| Average B factors (Å²) |  |
| Overall | 46.0 |
| Protein† | 43.1 |
| Tcc-Thr-Asp† | 39.0 |
| Ca²⁺† | 33.9 |
| Water† | 49.5 |
| Geometry |  |
| RMSD bond lengths (Å) | 0.014 |
| RMSD bond angles (°) | 0.624 |
| Bond-length RMSZ, all atoms | 0.19 |
| Bond-angle RMSZ, all atoms | 0.40 |
| MolProbity / validation |  |
| Ramachandran favored (%) | 98 |
| Ramachandran allowed (%) | 2 |
| Ramachandran outliers (%) | 0 |
| Rotamer outliers (%) | 1 |
| Clashscore | 1 |
| Cβ deviations (%) | 0 |
| Chirality outliers | 0 |
| Bond-length outliers | 0 |
| Bond-angle outliers | 0 |
| Ligand validation |  |
| Tcc residue RSCC | 0.94 |
| Tcc residue RSR | 0.09 |
| Tcc bond-length RMSZ | 0.44 |
| Tcc bond-angle RMSZ | 0.67 |
| Tcc bond-length outliers | 0 |
| Tcc bond-angle outliers | 0 |
| Tcc chirality outliers | 0 |
| Additional crystallographic statistics |  |
| Wilson B factor (Å²) | 36.6 |
| Twinning | None detected |
| Translational NCS | None detected |
| PDB accession | 37YK |

Values in parentheses correspond to the highest-resolution shell, 2.09–2.02 Å. †Component-specific B factors are taken from the final PHENIX refinement log; the wwPDB validation report gives an overall average B factor of 46.0 Å².

**Table S3. ImageJ-based hemagglutination inhibition analysis**

Raw ImageJ measurements, derived doughnut-to-pellet ratios, replicate-specific normalization, triplicate concentration-response summaries, and four-parameter variable-slope nonlinear-regression results are reported below. Whole-well and pellet regions of interest (ROIs) were quantified by ImageJ. Doughnut area = whole-well area − pellet area; doughnut mean gray value = [(whole area × whole mean) − (pellet area × pellet mean)]/doughnut area; raw ratio = doughnut mean/pellet mean. For each replicate, normalized hemagglutination was calculated using the corresponding WT and ΔFrhA-PBD mean raw ratios, with WT = 1 and ΔFrhA-PBD = 0. Values outside 0–1 were retained rather than truncated.

**Table S3A. Normalization controls.**

| Condition | Rep. | Whole area | Whole mean | Pellet area | Pellet mean | Doughnut area | Doughnut mean | Raw ratio |
| --- | --- | --- | --- | --- | --- | --- | --- | --- |
| WT | 1 | 0.0370 | 49953.5 | 0.0050 | 51044.4 | 0.0320 | 49783.1 | 0.9753 |
| WT | 2 | 0.0350 | 50484.3 | 0.0050 | 52895.9 | 0.0300 | 50082.4 | 0.9468 |
| WT | 3 | 0.0280 | 123.22 | 0.0040 | 109.94 | 0.0240 | 125.43 | 1.1408 |
| WT | 1 | 0.0370 | 49554.8 | 0.0050 | 50770.9 | 0.0320 | 49364.7 | 0.9723 |
| WT | 2 | 0.0350 | 49792.9 | 0.0050 | 52564.4 | 0.0300 | 49331.0 | 0.9385 |
| WT | 3 | 0.0280 | 124.31 | 0.0040 | 112 | 0.0240 | 126.36 | 1.1282 |
| WT | 1 | 0.0370 | 49551.0 | 0.0050 | 51028.4 | 0.0320 | 49320.2 | 0.9665 |
| WT | 2 | 0.0350 | 49773.1 | 0.0050 | 52946.6 | 0.0300 | 49244.1 | 0.9301 |
| WT | 3 | 0.0280 | 124.69 | 0.0040 | 113.24 | 0.0240 | 126.60 | 1.1180 |
| WT | 1 | 0.0370 | 49648.1 | 0.0050 | 51050.5 | 0.0320 | 49429.0 | 0.9682 |
| WT | 2 | 0.0350 | 49879.8 | 0.0050 | 53202.2 | 0.0300 | 49326.1 | 0.9271 |
| WT | 3 | 0.0320 | 126.10 | 0.0040 | 111.62 | 0.0280 | 128.17 | 1.1483 |
| WT | 1 | 0.0370 | 50013.3 | 0.0050 | 51413.8 | 0.0320 | 49794.5 | 0.9685 |
| WT | 2 | 0.0350 | 50051.2 | 0.0050 | 51566.0 | 0.0300 | 49798.7 | 0.9657 |
| WT | 3 | 0.0320 | 124.84 | 0.0040 | 110.79 | 0.0280 | 126.84 | 1.1449 |
| WT | 1 | 0.0370 | 50605.3 | 0.0050 | 51667.6 | 0.0320 | 50439.4 | 0.9762 |
| WT | 2 | 0.0310 | 49989.4 | 0.0050 | 51484.9 | 0.0260 | 49701.8 | 0.9654 |
| WT | 3 | 0.0320 | 122.89 | 0.0040 | 113.12 | 0.0280 | 124.29 | 1.0987 |
| WT | 1 | 0.0370 | 50345.3 | 0.0050 | 52303.0 | 0.0320 | 50039.4 | 0.9567 |
| WT | 2 | 0.0340 | 50224.9 | 0.0050 | 51833.2 | 0.0290 | 49947.7 | 0.9636 |
| WT | 3 | 0.0320 | 125.75 | 0.0040 | 113.98 | 0.0280 | 127.43 | 1.1181 |
| WT | 1 | 0.0370 | 51260.5 | 0.0050 | 51870.2 | 0.0320 | 51165.3 | 0.9864 |
| WT | 2 | 0.0340 | 50341.5 | 0.0050 | 52059.0 | 0.0290 | 50045.4 | 0.9613 |
| WT | 3 | 0.0320 | 113.92 | 0.0040 | 102.92 | 0.0280 | 115.49 | 1.1221 |
| ΔFrhA-PBD | 1 | 0.0370 | 46679.5 | 0.0050 | 57634.8 | 0.0320 | 44967.7 | 0.7802 |
| ΔFrhA-PBD | 2 | 0.0340 | 47082.3 | 0.0050 | 57472.5 | 0.0290 | 45290.9 | 0.7880 |
| ΔFrhA-PBD | 3 | 0.0320 | 160.59 | 0.0040 | 50.089 | 0.0280 | 176.37 | 3.5212 |
| ΔFrhA-PBD | 1 | 0.0370 | 46391.8 | 0.0050 | 57676.9 | 0.0320 | 44628.6 | 0.7738 |
| ΔFrhA-PBD | 2 | 0.0340 | 46580.6 | 0.0050 | 57221.0 | 0.0290 | 44746.0 | 0.7820 |
| ΔFrhA-PBD | 3 | 0.0320 | 158.95 | 0.0040 | 58.979 | 0.0280 | 173.23 | 2.9371 |
| ΔFrhA-PBD | 1 | 0.0370 | 46556.9 | 0.0040 | 57839.1 | 0.0330 | 45189.4 | 0.7813 |
| ΔFrhA-PBD | 2 | 0.0340 | 46508.8 | 0.0050 | 57265.8 | 0.0290 | 44654.1 | 0.7798 |
| ΔFrhA-PBD | 3 | 0.0320 | 158.53 | 0.0040 | 50.068 | 0.0280 | 174.02 | 3.4757 |
| ΔFrhA-PBD | 1 | 0.0370 | 46619.8 | 0.0040 | 58086.6 | 0.0330 | 45229.8 | 0.7787 |
| ΔFrhA-PBD | 2 | 0.0340 | 46703.1 | 0.0050 | 57414.9 | 0.0290 | 44856.2 | 0.7813 |
| ΔFrhA-PBD | 3 | 0.0320 | 154.41 | 0.0040 | 51.354 | 0.0280 | 169.14 | 3.2935 |
| ΔFrhA-PBD | 1 | 0.0370 | 47082.6 | 0.0030 | 58004.1 | 0.0340 | 46118.9 | 0.7951 |
| ΔFrhA-PBD | 2 | 0.0340 | 46728.1 | 0.0050 | 57105.3 | 0.0290 | 44938.9 | 0.7869 |
| ΔFrhA-PBD | 3 | 0.0320 | 152.18 | 0.0040 | 47.276 | 0.0280 | 167.17 | 3.5360 |
| ΔFrhA-PBD | 1 | 0.0370 | 46583 | 0.0040 | 58449.5 | 0.0330 | 45144.6 | 0.7724 |
| ΔFrhA-PBD | 2 | 0.0340 | 46849.0 | 0.0050 | 57118.5 | 0.0290 | 45078.4 | 0.7892 |
| ΔFrhA-PBD | 3 | 0.0320 | 152.14 | 0.0040 | 48.159 | 0.0280 | 167.00 | 3.4676 |
| ΔFrhA-PBD | 1 | 0.0370 | 47180.0 | 0.0040 | 58574.2 | 0.0330 | 45798.9 | 0.7819 |
| ΔFrhA-PBD | 2 | 0.0340 | 47251.8 | 0.0050 | 57469.5 | 0.0290 | 45490.1 | 0.7916 |
| ΔFrhA-PBD | 3 | 0.0320 | 149.49 | 0.0040 | 50.089 | 0.0280 | 163.69 | 3.2680 |
| ΔFrhA-PBD | 1 | 0.0370 | 47665.0 | 0.0040 | 58619.0 | 0.0330 | 46337.3 | 0.7905 |
| ΔFrhA-PBD | 2 | 0.0340 | 47919.0 | 0.0050 | 57886.9 | 0.0290 | 46200.4 | 0.7981 |
| ΔFrhA-PBD | 3 | 0.0320 | 142.79 | 0.0040 | 46.560 | 0.0280 | 156.53 | 3.3620 |

**Table S3B. Raw ImageJ measurements and derived normalized hemagglutination for inhibitor-treated wells.**

All individual inhibitor measurements used in the aggregate analysis are shown. *Normalized HA* is the replicate-specific value with WT = 1 and ΔFrhA-PBD = 0.

| Compound | Conc. (µM) | Rep. | Whole area | Whole mean | Pellet area | Pellet mean | Doughnut area | Doughnut mean | Raw ratio | Normalized HA | Inhibition (%) |
| --- | --- | --- | --- | --- | --- | --- | --- | --- | --- | --- | --- |
| L-10 | 1000 | 1 | 0.0370 | 49078.3 | 0.0060 | 55967.6 | 0.0310 | 47744.9 | 0.8531 | 0.3765 | 62.354 |
| L-10 | 1000 | 2 | 0.0340 | 49621.8 | 0.0050 | 55847.8 | 0.0290 | 48548.3 | 0.8693 | 0.5051 | 49.489 |
| L-10 | 1000 | 3 | 0.0320 | 142.51 | 0.0030 | 47.309 | 0.0290 | 152.36 | 3.2204 | 0.0615 | 93.848 |
| L-10 | 500 | 1 | 0.0370 | 48595.6 | 0.0060 | 56187.1 | 0.0310 | 47126.3 | 0.8387 | 0.3008 | 69.921 |
| L-10 | 500 | 2 | 0.0340 | 48987.1 | 0.0050 | 56785.3 | 0.0290 | 47642.5 | 0.8390 | 0.3189 | 68.112 |
| L-10 | 500 | 3 | 0.0320 | 135.17 | 0.0030 | 56.258 | 0.0290 | 143.33 | 2.5478 | 0.3631 | 63.688 |
| L-10 | 250 | 1 | 0.0370 | 48949.1 | 0.0040 | 56191.4 | 0.0330 | 48071.2 | 0.8555 | 0.3892 | 61.083 |
| L-10 | 250 | 2 | 0.0340 | 49233.3 | 0.0050 | 56279.4 | 0.0290 | 48018.4 | 0.8532 | 0.4063 | 59.373 |
| L-10 | 250 | 3 | 0.0320 | 131.04 | 0.0030 | 70.508 | 0.0290 | 137.30 | 1.9474 | 0.6323 | 36.766 |
| L-10 | 125 | 1 | 0.0370 | 49732.0 | 0.0040 | 55226.7 | 0.0330 | 49066.0 | 0.8884 | 0.5630 | 43.697 |
| L-10 | 125 | 2 | 0.0340 | 49812.5 | 0.0050 | 55035.6 | 0.0290 | 48912.0 | 0.8887 | 0.6246 | 37.543 |
| L-10 | 125 | 3 | 0.0320 | 127.69 | 0.0040 | 93.819 | 0.0280 | 132.53 | 1.4126 | 0.8721 | 12.787 |
| L-10 | 62.500 | 1 | 0.0370 | 50063.2 | 0.0007 | 55486.1 | 0.0363 | 49952.0 | 0.9003 | 0.6254 | 37.464 |
| L-10 | 62.500 | 2 | 0.0340 | 49968.6 | 0.0050 | 53226.0 | 0.0290 | 49407.0 | 0.9283 | 0.8674 | 13.256 |
| L-10 | 62.500 | 3 | 0.0320 | 126.34 | 0.0040 | 102.87 | 0.0280 | 129.69 | 1.2607 | 0.9402 | 5.9774 |
| L-10 | 31.200 | 1 | 0.0370 | 50010.4 | 0.0040 | 51942.9 | 0.0330 | 49776.1 | 0.9583 | 0.9315 | 6.8540 |
| L-10 | 31.200 | 2 | 0.0340 | 50221.7 | 0.0050 | 52100.7 | 0.0290 | 49897.7 | 0.9577 | 1.0485 | -4.8538 |
| L-10 | 31.200 | 3 | 0.0320 | 122.62 | 0.0040 | 108.26 | 0.0280 | 124.67 | 1.1516 | 0.9891 | 1.0875 |
| L-10 | 15.600 | 1 | 0.0370 | 50729.1 | 0.0040 | 52115.2 | 0.0330 | 50561.1 | 0.9702 | 0.9942 | 0.5790 |
| L-10 | 15.600 | 2 | 0.0340 | 50404.8 | 0.0050 | 51850.6 | 0.0290 | 50155.5 | 0.9673 | 1.1075 | -10.749 |
| L-10 | 15.600 | 3 | 0.0320 | 120.72 | 0.0040 | 107.72 | 0.0280 | 122.57 | 1.1379 | 0.9953 | 0.4733 |
| L-10 | 7.8000 | 1 | 0.0370 | 51169.4 | 0.0040 | 52551.6 | 0.0330 | 51001.9 | 0.9705 | 0.9960 | 0.4045 |
| L-10 | 7.8000 | 2 | 0.0340 | 50543.3 | 0.0050 | 51618.5 | 0.0290 | 50358.0 | 0.9756 | 1.1583 | -15.833 |
| L-10 | 7.8000 | 3 | 0.0320 | 117.56 | 0.0040 | 106.31 | 0.0280 | 119.17 | 1.1210 | 1.0029 | -0.2855 |
| D-10 | 1000 | 1 | 0.0370 | 49391.3 | 0.0040 | 55942.3 | 0.0330 | 48597.3 | 0.8687 | 0.4589 | 54.113 |
| D-10 | 1000 | 2 | 0.0340 | 49197.6 | 0.0050 | 55091.4 | 0.0290 | 48181.4 | 0.8746 | 0.5375 | 46.247 |
| D-10 | 1000 | 3 | 0.0320 | 147.13 | 0.0040 | 48.754 | 0.0280 | 161.19 | 3.3061 | 0.0231 | 97.690 |
| D-10 | 500 | 1 | 0.0370 | 48443.1 | 0.0040 | 56648.9 | 0.0330 | 47448.4 | 0.8376 | 0.2947 | 70.529 |
| D-10 | 500 | 2 | 0.0340 | 48910.8 | 0.0050 | 56616.7 | 0.0290 | 47582.2 | 0.8404 | 0.3277 | 67.232 |
| D-10 | 500 | 3 | 0.0320 | 140.84 | 0.0040 | 55.109 | 0.0280 | 153.08 | 2.7779 | 0.2600 | 74.004 |
| D-10 | 250 | 1 | 0.0370 | 48399.7 | 0.0040 | 56558.4 | 0.0330 | 47410.8 | 0.8383 | 0.2983 | 70.173 |
| D-10 | 250 | 2 | 0.0340 | 48685.0 | 0.0050 | 56520.8 | 0.0290 | 47334.0 | 0.8375 | 0.3095 | 69.054 |
| D-10 | 250 | 3 | 0.0320 | 139.28 | 0.0040 | 55.255 | 0.0280 | 151.28 | 2.7379 | 0.2779 | 72.211 |
| D-10 | 125 | 1 | 0.0370 | 48910.9 | 0.0040 | 56817.4 | 0.0330 | 47952.6 | 0.8440 | 0.3284 | 67.158 |
| D-10 | 125 | 2 | 0.0340 | 49001.2 | 0.0050 | 56722.8 | 0.0290 | 47669.9 | 0.8404 | 0.3275 | 67.248 |
| D-10 | 125 | 3 | 0.0320 | 138.51 | 0.0040 | 61.387 | 0.0280 | 149.53 | 2.4358 | 0.4133 | 58.666 |
| D-10 | 62.500 | 1 | 0.0370 | 48766.0 | 0.0040 | 56551.7 | 0.0330 | 47822.3 | 0.8456 | 0.3372 | 66.282 |
| D-10 | 62.500 | 2 | 0.0340 | 48756.5 | 0.0050 | 56211.2 | 0.0290 | 47471.2 | 0.8445 | 0.3528 | 64.720 |
| D-10 | 62.500 | 3 | 0.0320 | 135.63 | 0.0040 | 54.762 | 0.0280 | 147.19 | 2.6877 | 0.3004 | 69.963 |
| D-10 | 31.200 | 1 | 0.0370 | 48743.6 | 0.0040 | 57606.5 | 0.0330 | 47669.3 | 0.8275 | 0.2415 | 75.851 |
| D-10 | 31.200 | 2 | 0.0340 | 49103.6 | 0.0040 | 56987.7 | 0.0300 | 48052.4 | 0.8432 | 0.3448 | 65.524 |
| D-10 | 31.200 | 3 | 0.0320 | 137.07 | 0.0070 | 96.611 | 0.0250 | 148.40 | 1.5360 | 0.8168 | 18.322 |
| D-10 | 15.600 | 1 | 0.0370 | 49750.4 | 0.0040 | 57393.1 | 0.0330 | 48824.0 | 0.8507 | 0.3639 | 63.614 |
| D-10 | 15.600 | 2 | 0.0340 | 50051.3 | 0.0040 | 56663.1 | 0.0300 | 49169.8 | 0.8678 | 0.4956 | 50.436 |
| D-10 | 15.600 | 3 | 0.0320 | 127.20 | 0.0070 | 78.942 | 0.0250 | 140.71 | 1.7824 | 0.7063 | 29.370 |
| D-10 | 7.8000 | 1 | 0.0370 | 50786.6 | 0.0120 | 53651.0 | 0.0250 | 49411.7 | 0.9210 | 0.7347 | 26.533 |
| D-10 | 7.8000 | 2 | 0.0340 | 50755.5 | 0.0040 | 53498.7 | 0.0300 | 50389.8 | 0.9419 | 0.9513 | 4.8748 |
| D-10 | 7.8000 | 3 | 0.0320 | 117.05 | 0.0110 | 90.738 | 0.0210 | 130.82 | 1.4418 | 0.8590 | 14.097 |
| S-11 | 1000 | 1 | 0.0370 | 49992.6 | 0.0100 | 52994.9 | 0.0270 | 48880.7 | 0.9224 | 0.7420 | 25.803 |
| S-11 | 1000 | 2 | 0.0340 | 49934.6 | 0.0100 | 52939.1 | 0.0240 | 48682.8 | 0.9196 | 0.8143 | 18.572 |
| S-11 | 1000 | 3 | 0.0320 | 133.88 | 0.0080 | 92.549 | 0.0240 | 147.66 | 1.5955 | 0.7901 | 20.988 |
| S-11 | 500 | 1 | 0.0370 | 49789.8 | 0.0100 | 53091.7 | 0.0270 | 48566.9 | 0.9148 | 0.7019 | 29.809 |
| S-11 | 500 | 2 | 0.0340 | 49728.9 | 0.0060 | 54711.7 | 0.0280 | 48661.1 | 0.8894 | 0.6287 | 37.128 |
| S-11 | 500 | 3 | 0.0320 | 133.84 | 0.0080 | 91.550 | 0.0240 | 147.93 | 1.6159 | 0.7810 | 21.903 |
| S-11 | 250 | 1 | 0.0370 | 49788.8 | 0.0100 | 51822.5 | 0.0270 | 49035.6 | 0.9462 | 0.8678 | 13.218 |
| S-11 | 250 | 2 | 0.0340 | 49970.3 | 0.0060 | 53200.5 | 0.0280 | 49278.1 | 0.9263 | 0.8553 | 14.472 |
| S-11 | 250 | 3 | 0.0320 | 132.24 | 0.0080 | 96.072 | 0.0240 | 144.30 | 1.5020 | 0.8320 | 16.797 |
| S-11 | 125 | 1 | 0.0370 | 50052.4 | 0.0050 | 51766.9 | 0.0320 | 49784.6 | 0.9617 | 0.9495 | 5.0486 |
| S-11 | 125 | 2 | 0.0340 | 50192.7 | 0.0060 | 52019.9 | 0.0280 | 49801.2 | 0.9573 | 1.0463 | -4.6277 |
| S-11 | 125 | 3 | 0.0320 | 122.92 | 0.0080 | 94.595 | 0.0240 | 132.36 | 1.3992 | 0.8781 | 12.189 |
| S-11 | 62.500 | 1 | 0.0370 | 50237.7 | 0.0050 | 51472.6 | 0.0320 | 50044.7 | 0.9723 | 1.0052 | -0.5178 |
| S-11 | 62.500 | 2 | 0.0340 | 50002.7 | 0.0060 | 51630.9 | 0.0280 | 49653.8 | 0.9617 | 1.0731 | -7.3066 |
| S-11 | 62.500 | 3 | 0.0320 | 122.68 | 0.0080 | 104.27 | 0.0240 | 128.81 | 1.2354 | 0.9516 | 4.8418 |
| S-11 | 31.200 | 1 | 0.0370 | 50098.7 | 0.0050 | 51484.3 | 0.0320 | 49882.2 | 0.9689 | 0.9874 | 1.2637 |
| S-11 | 31.200 | 2 | 0.0340 | 50254.5 | 0.0060 | 51521.2 | 0.0280 | 49983.0 | 0.9701 | 1.1249 | -12.492 |
| S-11 | 31.200 | 3 | 0.0320 | 121.10 | 0.0080 | 105.70 | 0.0240 | 126.23 | 1.1942 | 0.9700 | 2.9985 |
| S-11 | 15.600 | 1 | 0.0370 | 50117.7 | 0.0050 | 51346.7 | 0.0320 | 49925.6 | 0.9723 | 1.0055 | -0.5528 |
| S-11 | 15.600 | 2 | 0.0340 | 50680.0 | 0.0060 | 52046.5 | 0.0280 | 50387.1 | 0.9681 | 1.1125 | -11.246 |
| S-11 | 15.600 | 3 | 0.0320 | 119.81 | 0.0080 | 108.71 | 0.0240 | 123.51 | 1.1362 | 0.9960 | 0.3960 |
| S-11 | 7.8000 | 1 | 0.0370 | 50501.5 | 0.0050 | 52052.1 | 0.0320 | 50259.2 | 0.9656 | 0.9698 | 3.0189 |
| S-11 | 7.8000 | 2 | 0.0340 | 50541.9 | 0.0060 | 51246.8 | 0.0280 | 50390.8 | 0.9833 | 1.2058 | -20.575 |
| S-11 | 7.8000 | 3 | 0.0320 | 113.46 | 0.0080 | 101.81 | 0.0240 | 117.34 | 1.1525 | 0.9887 | 1.1273 |
| R-11 | 1000 | 1 | 0.0370 | 50592.8 | 0.0130 | 51996.7 | 0.0240 | 49832.4 | 0.9584 | 0.9319 | 6.8061 |
| R-11 | 1000 | 2 | 0.0340 | 50206.2 | 0.0120 | 50681.3 | 0.0220 | 49947.1 | 0.9855 | 1.2194 | -21.937 |
| R-11 | 1000 | 3 | 0.0320 | 123.44 | 0.0080 | 106.06 | 0.0240 | 129.23 | 1.2185 | 0.9591 | 4.0870 |
| R-11 | 500 | 1 | 0.0370 | 48465.5 | 0.0070 | 54621.0 | 0.0300 | 47029.2 | 0.8610 | 0.4183 | 58.172 |
| R-11 | 500 | 2 | 0.0340 | 49422.4 | 0.0120 | 52614.3 | 0.0220 | 47681.4 | 0.9062 | 0.7322 | 26.781 |
| R-11 | 500 | 3 | 0.0320 | 147.19 | 0.0050 | 54.964 | 0.0270 | 164.27 | 2.9886 | 0.1655 | 83.455 |
| R-11 | 250 | 1 | 0.0370 | 48120.9 | 0.0050 | 56150.5 | 0.0320 | 46866.3 | 0.8347 | 0.2792 | 72.076 |
| R-11 | 250 | 2 | 0.0340 | 48934.4 | 0.0060 | 55215.8 | 0.0280 | 47588.3 | 0.8619 | 0.4594 | 54.059 |
| R-11 | 250 | 3 | 0.0320 | 145.21 | 0.0050 | 55.917 | 0.0270 | 161.74 | 2.8925 | 0.2086 | 79.144 |
| R-11 | 125 | 1 | 0.0370 | 49221.1 | 0.0050 | 56545.9 | 0.0320 | 48076.6 | 0.8502 | 0.3614 | 63.863 |
| R-11 | 125 | 2 | 0.0340 | 48358.2 | 0.0040 | 56485.2 | 0.0300 | 47274.6 | 0.8369 | 0.3062 | 69.376 |
| R-11 | 125 | 3 | 0.0320 | 139.12 | 0.0050 | 61.639 | 0.0270 | 153.46 | 2.4897 | 0.3892 | 61.083 |
| R-11 | 62.500 | 1 | 0.0370 | 49636.7 | 0.0050 | 56367.8 | 0.0320 | 48585.0 | 0.8619 | 0.4231 | 57.688 |
| R-11 | 62.500 | 2 | 0.0340 | 49303.9 | 0.0040 | 56470.3 | 0.0300 | 48348.3 | 0.8562 | 0.4245 | 57.554 |
| R-11 | 62.500 | 3 | 0.0320 | 132.30 | 0.0050 | 56.722 | 0.0270 | 146.30 | 2.5793 | 0.3490 | 65.099 |
| R-11 | 31.200 | 1 | 0.0370 | 49583.2 | 0.0100 | 52387.9 | 0.0270 | 48544.4 | 0.9266 | 0.7645 | 23.552 |
| R-11 | 31.200 | 2 | 0.0340 | 49581.8 | 0.0060 | 54317.8 | 0.0280 | 48567.0 | 0.8941 | 0.6577 | 34.228 |
| R-11 | 31.200 | 3 | 0.0320 | 125.06 | 0.0070 | 81.611 | 0.0250 | 137.23 | 1.6815 | 0.7516 | 24.844 |
| R-11 | 15.600 | 1 | 0.0370 | 50089.7 | 0.0060 | 52111.5 | 0.0310 | 49698.4 | 0.9537 | 0.9072 | 9.2763 |
| R-11 | 15.600 | 2 | 0.0340 | 50570.9 | 0.0060 | 54116.2 | 0.0280 | 49811.2 | 0.9204 | 0.8195 | 18.051 |
| R-11 | 15.600 | 3 | 0.0320 | 119.67 | 0.0070 | 98.287 | 0.0250 | 125.66 | 1.2785 | 0.9323 | 6.7743 |
| R-11 | 7.8000 | 1 | 0.0370 | 51144.9 | 0.0060 | 52271.3 | 0.0310 | 50926.9 | 0.9743 | 1.0158 | -1.5847 |
| R-11 | 7.8000 | 2 | 0.0340 | 50582.1 | 0.0050 | 52327.8 | 0.0290 | 50281.1 | 0.9609 | 1.0680 | -6.8028 |
| R-11 | 7.8000 | 3 | 0.0320 | 116.21 | 0.0070 | 103.35 | 0.0250 | 119.81 | 1.1592 | 0.9857 | 1.4273 |

**Table S3C. Triplicate normalized concentration-response data used for curve fitting.**

| Compound | Conc. (µM) | n | Mean normalized HA | SD normalized HA | Mean inhibition (%) | SD inhibition (%) |
| --- | --- | --- | --- | --- | --- | --- |
| L-10 | 1000 | 3 | 0.3144 | 0.2282 | 68.564 | 22.822 |
| L-10 | 500 | 3 | 0.3276 | 0.0321 | 67.240 | 3.2068 |
| L-10 | 250 | 3 | 0.4759 | 0.1357 | 52.407 | 13.573 |
| L-10 | 125 | 3 | 0.6866 | 0.1636 | 31.342 | 16.361 |
| L-10 | 62.500 | 3 | 0.8110 | 0.1648 | 18.899 | 16.485 |
| L-10 | 31.200 | 3 | 0.9897 | 0.0585 | 1.0292 | 5.8541 |
| L-10 | 15.600 | 3 | 1.0323 | 0.0651 | -3.2324 | 6.5102 |
| L-10 | 7.8000 | 3 | 1.0524 | 0.0918 | -5.2379 | 9.1818 |
| D-10 | 1000 | 3 | 0.3398 | 0.2771 | 66.017 | 27.711 |
| D-10 | 500 | 3 | 0.2941 | 0.0339 | 70.588 | 3.3862 |
| D-10 | 250 | 3 | 0.2952 | 0.0160 | 70.479 | 1.6006 |
| D-10 | 125 | 3 | 0.3564 | 0.0493 | 64.357 | 4.9293 |
| D-10 | 62.500 | 3 | 0.3301 | 0.0269 | 66.988 | 2.6921 |
| D-10 | 31.200 | 3 | 0.4677 | 0.3067 | 53.232 | 30.671 |
| D-10 | 15.600 | 3 | 0.5219 | 0.1727 | 47.807 | 17.273 |
| D-10 | 7.8000 | 3 | 0.8483 | 0.1087 | 15.168 | 10.869 |
| S-11 | 1000 | 3 | 0.7821 | 0.0368 | 21.788 | 3.6812 |
| S-11 | 500 | 3 | 0.7039 | 0.0761 | 29.613 | 7.6143 |
| S-11 | 250 | 3 | 0.8517 | 0.0182 | 14.829 | 1.8159 |
| S-11 | 125 | 3 | 0.9580 | 0.0844 | 4.2034 | 8.4404 |
| S-11 | 62.500 | 3 | 1.0099 | 0.0609 | -0.9942 | 6.0882 |
| S-11 | 31.200 | 3 | 1.0274 | 0.0849 | -2.7432 | 8.4870 |
| S-11 | 15.600 | 3 | 1.0380 | 0.0647 | -3.8010 | 6.4652 |
| S-11 | 7.8000 | 3 | 1.0548 | 0.1311 | -5.4764 | 13.110 |
| R-11 | 1000 | 0 | — | — | — | — |
| R-11 | 500 | 3 | 0.4386 | 0.2839 | 56.136 | 28.392 |
| R-11 | 250 | 3 | 0.3157 | 0.1293 | 68.426 | 12.934 |
| R-11 | 125 | 3 | 0.3523 | 0.0422 | 64.774 | 4.2210 |
| R-11 | 62.500 | 3 | 0.3989 | 0.0432 | 60.114 | 4.3178 |
| R-11 | 31.200 | 3 | 0.7246 | 0.0583 | 27.541 | 5.8270 |
| R-11 | 15.600 | 3 | 0.8863 | 0.0592 | 11.367 | 5.9220 |
| R-11 | 7.8000 | 3 | 1.0232 | 0.0416 | -2.3201 | 4.1640 |
| L-10 | 1 | 8 | 0.9044 | 0.0550 | — | — |
| L-10 | 2 | 8 | 0.9099 | 0.0542 | — | — |
| L-10 | 3 | 8 | 1.7249 | 0.7851 | — | — |
| D-10 | 1 | 8 | 0.8542 | 0.0295 | — | — |
| D-10 | 2 | 8 | 0.8613 | 0.0354 | — | — |
| D-10 | 3 | 8 | 2.3382 | 0.6738 | — | — |
| S-11 | 1 | 8 | 0.9530 | 0.0229 | — | — |
| S-11 | 2 | 8 | 0.9470 | 0.0319 | — | — |
| S-11 | 3 | 8 | 1.3539 | 0.1994 | — | — |
| R-11 | 1 | 8 | 0.9026 | 0.0563 | — | — |
| R-11 | 2 | 8 | 0.9028 | 0.0518 | — | — |
| R-11 | 3 | 8 | 2.0360 | 0.7818 | — | — |

**Table S3D. Four-parameter variable-slope nonlinear-regression results.**

Model: $Y = Bottom +\frac{\left( Top - Bottom \right)}{\left[ 1 + \left( \frac{X}{IC50} \right)^{HillSlope} \right]}$. The normalized vehicle control was included at X = 0 µM and Y = 100%. Fits used the mean of the three normalized replicate values at each inhibitor concentration. The 95% confidence interval was calculated from the standard error of log10(IC50) from the nonlinear least-squares covariance matrix using a two-sided t critical value with df = number of fitted concentration points − 4.

| Compound | IC50 (µM) | Regression SE of IC50 (µM) | 95% CI (µM) | Bottom | Top | Hill slope | log10(IC50) | R² | Fit points | Max included (µM) |
| --- | --- | --- | --- | --- | --- | --- | --- | --- | --- | --- |
| L-10 | 106.20 | 14.120 | 75.45–149.47 | -6.1471 | 107.12 | 1.8094 | 2.0261 | 0.9882 | 9 | 1000 |
| D-10 | 11.077 | 1.0616 | 8.66–14.17 | -10.248 | 100.49 | 3.3399 | 1.0444 | 0.9824 | 9 | 1000 |
| S-11 | 191.60 | 39.516 | 112.75–325.57 | -20.450 | 115.35 | 3.0872 | 2.2824 | 0.9415 | 9 | 1000 |
| R-11 | 27.939 | 6.6373 | 14.45–54.03 | -31.482 | 101.90 | 2.9911 | 1.4462 | 0.9347 | 8 | 500 |

**Table S4. Complete data and nonlinear regression analysis of the V. cholerae biofilm inhibition assay.**

Biofilm formation by WT V. cholerae O395 was measured as A570 after crystal-violet staining. Raw absorbance values, controls used for normalization, normalized values, and IC50 curve-fitting results are reported below. Compound identifiers follow the manuscript nomenclature: 1, D-**10**, and R-**11**.

**A. Raw inhibitor data (A570)**

| Concentration (µM) | 1 Rep 1 | 1 Rep 2 | 1 Rep 3 | D-10 Rep 1 | D-10 Rep 2 | D-10 Rep 3 | R-11 Rep 1 | R-11 Rep 2 | R-11 Rep 3 |
| --- | --- | --- | --- | --- | --- | --- | --- | --- | --- |
| 125 | 0.392 | 0.355 | 0.282 | 0.269 | 0.251 | 0.277 | 0.255 | 0.271 | 0.265 |
| 62.5 | 0.468 | 0.309 | 0.344 | 0.276 | 0.306 | 0.351 | 0.292 | 0.299 | 0.271 |
| 31.25 | 0.440 | 0.444 | 0.413 | 0.363 | 0.256 | 0.293 | 0.304 | 0.308 | 0.283 |
| 15.625 | 0.652 | 0.507 | 0.535 | 0.343 | 0.276 | 0.340 | 0.305 | 0.268 | 0.309 |
| 7.815 | 0.610 | 0.596 | 0.645 | 0.360 | 0.378 | 0.438 | 0.319 | 0.292 | 0.346 |
| 3.9075 | 0.684 | 0.664 | 0.645 | 0.462 | 0.321 | 0.469 | 0.358 | 0.359 | 0.344 |
| 1.95375 | 0.587 | 0.726 | 0.638 | 0.504 | 0.534 | 0.518 | 0.576 | 0.600 | 0.522 |
| 0.976875 | 0.752 | 0.688 | 0.588 | 0.628 | 0.702 | 0.593 | 0.862 | 0.788 | 0.856 |

**B. Controls used for normalization (A570)**

| Control | Rep 1 | Rep 2 | Rep 3 | Mean | SD |
| --- | --- | --- | --- | --- | --- |
| Vehicle (DMSO) | 0.092 | 0.095 | 0.092 | 0.093 | 0.002 |
| WT V. cholerae | 0.931 | 0.811 | 0.827 | 0.856 | 0.065 |
| ΔFrhA-PBD V. cholerae | 0.208 | 0.191 | 0.228 | 0.209 | 0.019 |

Normalization used the mean WT control (A570 = 0.856) as 100% biofilm and the mean ΔFrhA-PBD control (A570 = 0.209) as 0% biofilm. The vehicle control (mean A570 = 0.093) is reported as a plate/background control and was not used as a normalization anchor. For each inhibitor measurement A, normalized biofilm (%) = 100 × (A − 0.209) / (0.856 − 0.209).

**C. Normalized biofilm data (%)**

| Concentration (µM) | 1 Rep 1 | 1 Rep 2 | 1 Rep 3 | D-10 Rep 1 | D-10 Rep 2 | D-10 Rep 3 | R-11 Rep 1 | R-11 Rep 2 | R-11 Rep 3 |
| --- | --- | --- | --- | --- | --- | --- | --- | --- | --- |
| 125 | 28.270 | 22.554 | 11.277 | 9.269 | 6.488 | 10.505 | 7.106 | 9.578 | 8.651 |
| 62.5 | 40.010 | 15.448 | 20.855 | 10.350 | 14.985 | 21.936 | 12.822 | 13.903 | 9.578 |
| 31.25 | 35.685 | 36.303 | 31.514 | 23.790 | 7.261 | 12.976 | 14.676 | 15.294 | 11.432 |
| 15.625 | 68.435 | 46.035 | 50.360 | 20.700 | 10.350 | 20.237 | 14.830 | 9.114 | 15.448 |
| 7.815 | 61.946 | 59.784 | 67.353 | 23.326 | 26.107 | 35.376 | 16.993 | 12.822 | 21.164 |
| 3.9075 | 73.378 | 70.288 | 67.353 | 39.083 | 17.302 | 40.165 | 23.018 | 23.172 | 20.855 |
| 1.95375 | 58.393 | 79.866 | 66.272 | 45.572 | 50.206 | 47.734 | 56.694 | 60.402 | 48.352 |
| 0.976875 | 83.883 | 73.996 | 58.548 | 64.727 | 76.159 | 59.320 | 100.875 | 89.444 | 99.949 |

**D. Nonlinear curve fitting and IC50 estimates**

| Compound | IC50 (µM) | | SE, IC50 (µM) | | 95% CI, IC50 (µM) | | Hill slope | | 95% CI, Hill slope |
| --- | --- | --- | --- | --- | --- | --- | --- | --- | --- |
| 1 | 13 | | 2.189 | | 8.3–17 | | −0.5810 | | −0.7246 to −0.4374 |
| D-10 | 1.6 | | 0.4007 | | 0.80–2.5 | | −0.5648 | | −0.6939 to −0.4357 |
| R-11 | 2.2 | | 0.5934 | | 0.95–3.4 | | −0.7516 | | −0.9767 to −0.5264 |
| Compound | | **df** | | **Weighted R²** | | **Weighted sum of squares (1/Y)** | | **Sy.x** | |
| 1 | | 22 | | 0.8236 | | 46.17 | | 1.449 | |
| D-10 | | 22 | | 0.8188 | | 49.69 | | 1.503 | |
| R-11 | | 22 | | 0.7431 | | 115.0 | | 2.287 | |

The revised Prism nonlinear-regression analysis constrained IC50 > 0, fitted IC50 and Hill slope separately for each data set, and used 24 Y values for each compound. The fits were weighted by **1**/Y and asymptotic 95% confidence intervals are reported. A global comparison of the three data sets rejected the model in which IC50 was shared among all compounds (F(2, 66) = 18.04, P < 0.0001), indicating that IC50 differed for at least one compound. Planned two-data-set global F tests were then used to compare compound **1** with D-**10** and R-**11**. IC50 values and their 95% confidence limits are reported to two significant figures.

**E. Pairwise comparison of IC50 fits**

For each planned comparison, Prism compared a model in which the two data sets shared a single IC50 while allowing separate Hill slopes with a model allowing separate IC50 values and separate Hill slopes. Both comparisons rejected the shared-IC50 model. The fitted IC50 values were lower for D-**10** and R-**11** than for compound 1, supporting significantly greater potency in this assay.

| Comparison | Separate-fit IC50 values (µM) | Shared-fit IC50 (µM) | F (DFn, DFd) | P value | Interpretation |
| --- | --- | --- | --- | --- | --- |
| 1 vs D-10 | **1**: 13; D-**10**: 1.6 | 2.913 | 51.45 (1, 44) | <0.0001 | D-**10** has a significantly lower IC50 than 1 |
| 1 vs R-11 | **1**: 13; R-**11**: 2.2 | 3.057 | 26.26 (1, 44) | <0.0001 | R-**11** has a significantly lower IC50 than 1 |

The fitted IC50 values and asymptotic 95% confidence intervals were: **1**, 13 µM (8.3–17 µM); D-**10**, 1.6 µM (0.80–2.5 µM); and R-**11**, 2.2 µM (0.95–3.4 µM).

The pairwise F tests directly tested equality of IC50 while allowing Hill slopes to differ; therefore, the significant tests support the conclusion that D-**10** and R-**11** have lower IC50 values than compound 1.

**Table S5. Caco 2 TccTD Dose Response Data Processing and Statistical Analysis.**

**Table S5A. Raw responses and row statistics**

| Concentration (µM) | Sample | Replicate 1 | Replicate 2 | Replicate 3 | Mean | SD | n |
| --- | --- | --- | --- | --- | --- | --- | --- |
| 1000.00000 | **L-1** | 0.335 | 0.444 | 0.377 | 0.385 | 0.055 | 3 |
| 1000.00000 | **D-10** | 0.188 | 0.176 | 0.129 | 0.164 | 0.031 | 3 |
| 1000.00000 | **R-11** | 0.191 | 0.214 | 0.192 | 0.199 | 0.013 | 3 |
| 1000.00000 | **S-11** | 0.285 | 0.204 | 0.179 | 0.223 | 0.055 | 3 |
| 333.35000 | **L-1** | 0.464 | 0.542 | 0.461 | 0.489 | 0.046 | 3 |
| 333.35000 | **D-10** | 0.209 | 0.268 | 0.247 | 0.241 | 0.030 | 3 |
| 333.35000 | **R-11** | 0.272 | 0.359 | 0.199 | 0.277 | 0.080 | 3 |
| 333.35000 | **S-11** | 0.496 | 0.775 | 0.583 | 0.618 | 0.143 | 3 |
| 111.10000 | **L-1** | 1.606 | 1.085 | 1.078 | 1.256 | 0.303 | 3 |
| 111.10000 | **D-10** | 0.457 | 0.481 | 0.733 | 0.557 | 0.153 | 3 |
| 111.10000 | **R-11** | 0.534 | 0.627 | 0.270 | 0.477 | 0.185 | 3 |
| 111.10000 | **S-11** | 1.410 | 1.368 | 1.157 | 1.312 | 0.136 | 3 |
| 37.03500 | **L-1** | 1.723 | 1.473 | 1.640 | 1.612 | 0.127 | 3 |
| 37.03500 | **D-10** | 0.870 | 0.924 | 1.008 | 0.934 | 0.070 | 3 |
| 37.03500 | **R-11** | 0.779 | 0.875 | 0.502 | 0.719 | 0.194 | 3 |
| 37.03500 | **S-11** | 1.389 | 1.575 | 1.489 | 1.484 | 0.093 | 3 |
| 12.34500 | **L-1** | 2.162 | 2.044 | 1.847 | 2.018 | 0.159 | 3 |
| 12.34500 | **D-10** | 1.568 | 1.433 | 1.402 | 1.468 | 0.088 | 3 |
| 12.34500 | **R-11** | 1.077 | 1.365 | 0.923 | 1.122 | 0.224 | 3 |
| 12.34500 | **S-11** | 1.846 | 1.766 | 1.649 | 1.754 | 0.099 | 3 |
| 4.11500 | **L-1** | 2.044 | 1.889 | 1.812 | 1.915 | 0.118 | 3 |
| 4.11500 | **D-10** | 1.407 | 1.417 | 1.609 | 1.478 | 0.114 | 3 |
| 4.11500 | **R-11** | 1.544 | 1.528 | 1.065 | 1.379 | 0.272 | 3 |
| 4.11500 | **S-11** | 2.039 | 1.683 | 1.630 | 1.784 | 0.222 | 3 |
| 1.37150 | **L-1** | 2.188 | 1.833 | 1.980 | 2.000 | 0.178 | 3 |
| 1.37150 | **D-10** | 1.894 | 1.686 | 1.949 | 1.843 | 0.139 | 3 |
| 1.37150 | **R-11** | 1.593 | 1.524 | 1.235 | 1.451 | 0.190 | 3 |
| 1.37150 | **S-11** | 1.752 | 1.704 | 1.623 | 1.693 | 0.065 | 3 |
| 0.45725 | **L-1** | 1.992 | 2.014 | 1.704 | 1.903 | 0.173 | 3 |
| 0.45725 | **D-10** | 1.840 | 1.940 | 2.035 | 1.938 | 0.098 | 3 |
| 0.45725 | **R-11** | 2.286 | 2.370 | 2.013 | 2.223 | 0.187 | 3 |
| 0.45725 | **S-11** | 1.834 | 1.691 | 1.483 | 1.669 | 0.177 | 3 |
| 0.15240 | **L-1** | 2.082 | 1.926 | 1.792 | 1.933 | 0.145 | 3 |
| 0.15240 | **D-10** | 1.943 | 1.724 | 1.850 | 1.839 | 0.110 | 3 |
| 0.15240 | **R-11** | 2.487 | 2.620 | 2.065 | 2.391 | 0.290 | 3 |
| 0.15240 | **S-11** | 1.760 | 1.537 | 1.538 | 1.612 | 0.128 | 3 |
| 0 | FrhASplit-PBD control | 1.745 | 1.632 | 1.557 | 1.645 | 0.095 | 3 |
| 0 | Vehicle control | 0.186 | 0.184 | 0.156 | 0.175 | 0.017 | 3 |

*Raw values, means, SDs, and replicate counts reproduce the Prism exports. FrhASplit-PBD control: 1.645 ± 0.095; vehicle control: 0.175 ± 0.017 (mean ± SD, n = 3).*

**Table S5B. Normalized response values and row statistics**

| Concentration (µM) | Compound | Replicate 1 (%) | Replicate 2 (%) | Replicate 3 (%) | Mean (%) | SD | n |
| --- | --- | --- | --- | --- | --- | --- | --- |
| 1000.000 | **L-1** | 10.887 | 18.303 | 13.745 | 14.312 | 3.740 | 3 |
| 1000.000 | **D-10** | 0.885 | 0.068 | -3.130 | -0.726 | 2.122 | 3 |
| 1000.000 | **R-11** | 1.089 | 2.654 | 1.157 | 1.633 | 0.885 | 3 |
| 1000.000 | **S-11** | 7.485 | 1.973 | 0.272 | 3.243 | 3.771 | 3 |
| 333.350 | **L-1** | 19.664 | 24.972 | 19.460 | 21.365 | 3.125 | 3 |
| 333.350 | **D-10** | 2.313 | 6.328 | 4.899 | 4.513 | 2.035 | 3 |
| 333.350 | **R-11** | 6.600 | 12.520 | 1.633 | 6.918 | 5.450 | 3 |
| 333.350 | **S-11** | 21.842 | 40.826 | 27.761 | 30.143 | 9.714 | 3 |
| 111.100 | **L-1** | 97.369 | 61.919 | 61.443 | 73.577 | 20.606 | 3 |
| 111.100 | **D-10** | 19.188 | 20.821 | 37.968 | 25.992 | 10.403 | 3 |
| 111.100 | **R-11** | 24.427 | 30.755 | 6.464 | 20.549 | 12.601 | 3 |
| 111.100 | **S-11** | 84.033 | 81.175 | 66.818 | 77.342 | 9.225 | 3 |
| 37.035 | **L-1** | 105.330 | 88.319 | 99.682 | 97.777 | 8.664 | 3 |
| 37.035 | **D-10** | 47.290 | 50.964 | 56.680 | 51.645 | 4.732 | 3 |
| 37.035 | **R-11** | 41.098 | 47.630 | 22.250 | 36.993 | 13.179 | 3 |
| 37.035 | **S-11** | 82.604 | 95.260 | 89.408 | 89.091 | 6.334 | 3 |
| 12.345 | **L-1** | 135.201 | 127.172 | 113.767 | 125.380 | 10.829 | 3 |
| 12.345 | **D-10** | 94.783 | 85.598 | 83.488 | 87.956 | 6.005 | 3 |
| 12.345 | **R-11** | 61.374 | 80.971 | 50.896 | 64.414 | 15.266 | 3 |
| 12.345 | **S-11** | 113.699 | 108.256 | 100.295 | 107.417 | 6.741 | 3 |
| 4.115 | **L-1** | 127.172 | 116.625 | 111.386 | 118.394 | 8.040 | 3 |
| 4.115 | **D-10** | 83.829 | 84.509 | 97.573 | 88.637 | 7.746 | 3 |
| 4.115 | **R-11** | 93.150 | 92.062 | 60.558 | 81.923 | 18.511 | 3 |
| 4.115 | **S-11** | 126.831 | 102.608 | 99.002 | 109.480 | 15.134 | 3 |
| 1.372 | **L-1** | 136.970 | 112.815 | 122.817 | 124.201 | 12.137 | 3 |
| 1.372 | **D-10** | 116.965 | 102.812 | 120.708 | 113.495 | 9.439 | 3 |
| 1.372 | **R-11** | 96.484 | 91.790 | 72.125 | 86.800 | 12.924 | 3 |
| 1.372 | **S-11** | 107.303 | 104.037 | 98.526 | 103.289 | 4.436 | 3 |
| 0.457 | **L-1** | 123.633 | 125.130 | 104.037 | 117.600 | 11.770 | 3 |
| 0.457 | **D-10** | 113.291 | 120.095 | 126.559 | 119.982 | 6.635 | 3 |
| 0.457 | **R-11** | 143.638 | 149.354 | 125.062 | 139.351 | 12.701 | 3 |
| 0.457 | **S-11** | 112.883 | 103.153 | 89.000 | 101.679 | 12.010 | 3 |
| 0.152 | **L-1** | 129.757 | 119.143 | 110.025 | 119.642 | 9.875 | 3 |
| 0.152 | **D-10** | 120.299 | 105.398 | 113.971 | 113.223 | 7.479 | 3 |
| 0.152 | **R-11** | 157.315 | 166.364 | 128.601 | 150.760 | 19.716 | 3 |
| 0.152 | **S-11** | 107.848 | 92.674 | 92.742 | 97.755 | 8.741 | 3 |

*Normalized replicate values reproduce the Prism export. Means and sample SDs were calculated from the displayed normalized values. Minor differences can arise when normalization is recalculated from the rounded raw CSV values.*

**Table S5C. Nonlinear regression parameter estimates and fit diagnostics**

**Parameter estimates**

| Compound | IC50 (µM) | IC50 95% CI (µM) | logIC50 | logIC50 95% CI | Constraint |
| --- | --- | --- | --- | --- | --- |
| L-1 | 222.8 | 126.6 to 403.0 | 2.348 | 2.103 to 2.605 | IC50 > 0 |
| D-10 | 44.34 | 31.04 to 63.44 | 1.647 | 1.492 to 1.802 | IC50 > 0 |
| R-11 | 23.59 | 11.01 to 51.22 | 1.373 | 1.042 to 1.709 | IC50 > 0 |
| S-11 | 211.2 | 151.8 to 294.8 | 2.325 | 2.181 to 2.469 | IC50 > 0 |

**Goodness of fit and number of observations**

| Compound | Degrees of freedom | R squared | Sum of squares | Sy.x | X values | Y values analyzed |
| --- | --- | --- | --- | --- | --- | --- |
| L-1 | 26 | 0.7586 | 11850 | 21.35 | 27 | 27 |
| D-10 | 26 | 0.9265 | 4124 | 12.59 | 27 | 27 |
| R-11 | 26 | 0.7778 | 16597 | 25.27 | 27 | 27 |
| S-11 | 26 | 0.8949 | 3755 | 12.02 | 27 | 27 |

*All 27 normalized values per compound were included in the nonlinear regression. Confidence intervals are profile-likelihood intervals.*

**Table S5D. Pairwise extra sum-of-squares F tests comparing fitted curves**

| Comparison | Separate IC50 values (µM) | Shared IC50 (µM) | Separate-model SS | Shared-model SS | F (DFn, DFd) | P value | Conclusion |
| --- | --- | --- | --- | --- | --- | --- | --- |
| L-1 vs D-10 | L-1 222.8; D-10 44.34 | 107.7 | 15974 | 23219 | 23.59 (1, 52) | <0.0001 | Reject shared-curve model |
| D-10 vs S-11 | D-10 44.34; S-11 211.2 | 100.3 | 7879 | 14399 | 43.03 (1, 52) | <0.0001 | Reject shared-curve model |
| L-1 vs R-11 | L-1 222.8; R-11 23.59 | 89.34 | 28447 | 41295 | 23.49 (1, 52) | <0.0001 | Reject shared-curve model |
| R-11 vs S-11 | R-11 23.59; S-11 211.2 | 80.87 | 20352 | 32139 | 30.11 (1, 52) | <0.0001 | Reject shared-curve model |

*Null model: one shared IC50. Alternative model: separate IC50 values. All comparisons rejected the null model at alpha = 0.05.*

**Table S6. One-phase exponential-decay analysis of chymotrypsin digestion of AGWTD and AGwTD.**

The Prism one-phase decay model was fitted with K constrained to K > 0. Compound nomenclature is standardized as L-**2** = H-Ala-Gly-Trp-Thr-Asp-OH (AGWTD) and D-**2** = H-Ala-Gly-D-Trp-Thr-Asp-OH (AGwTD).

| Parameter | D-2 (AGwTD; summed peaks) | L-2 (AGWTD) | L-2 fragment (AGW) |
| --- | --- | --- | --- |
| Y0 | 100.0 | 96.53 | 2.116 |
| Plateau | 88.60 | 2.008 | 99.45 |
| K | Unstable | 0.03248 | 0.03420 |
| Half-life (min) | Unstable | 21.34 | 20.27 |
| Tau (min) | Unstable | 30.79 | 29.24 |
| Span | 11.40 | 94.53 | -97.34 |
| 95% CI, Y0 | 81.19 to 118.8 | 89.31 to 104.0 | -3.182 to 7.318 |
| 95% CI, Plateau | 82.65 to 94.55 | -8.917 to 10.82 | 93.24 to 106.6 |
| 95% CI, K | Very wide | 0.02331 to 0.04380 | 0.02744 to 0.04205 |
| 95% CI, half-life (min) | Very wide | 15.82 to 29.73 | 16.48 to 25.26 |
| 95% CI, Tau (min) | Very wide | 22.83 to 42.90 | 23.78 to 36.44 |
| Degrees of freedom | 8 | 9 | 9 |
| R² | 0.1816 | 0.9814 | 0.9912 |
| Sum of squares | 532.4 | 244.5 | 122.5 |
| Sy.x | 8.158 | 5.213 | 3.690 |
| Constraint on K | K > 0 | K > 0 | K > 0 |
| # X values | 24 | 24 | 24 |
| # Y values analyzed | 11 | 12 | 12 |

**Table S7. One-phase exponential-decay analysis of chymotrypsin digestion of peptides and Tcc analogs.**

The Prism one-phase decay model was fitted independently to each compound with K constrained to K > 0.

| Parameter | D-3 | L-3 | D-10 | L-10 | *R*-11 | *S*-11 | L-1 |
| --- | --- | --- | --- | --- | --- | --- | --- |
| Y0 | 104.9 | 99.20 | 100.0 | 99.76 | 97.38 | 102.2 | 113.6 |
| Plateau | 103.8 | 108.2 | 105.1 | 105.6 | 105.1 | Unstable | 15.79 |
| K | 0.03328 | 0.08665 | 0.5243 | 0.1559 | 0.05734 | 3.244e-005 | 0.02103 |
| Half-life (min) | 20.83 | 8.000 | 1.322 | 4.446 | 12.09 | 21364 | 32.96 |
| Tau (min) | 30.05 | 11.54 | 1.907 | 6.414 | 17.44 | 30822 | 47.56 |
| Span | 1.036 | -8.969 | -5.126 | -5.819 | -7.729 | Unstable | 97.81 |
| 95% CI, Y0 | 80.30 to 121.2 | 87.78 to 110.5 | 93.45 to 106.5 | 95.79 to 103.7 | 74.04 to 124.4 | 36.14 to 137.3 | 66.55 to 170.1 |
| 95% CI, Plateau | 96.39 to 114.0 | 101.7 to 111.1 | 102.2 to 108.0 | 103.3 to 108.0 | 90.55 to 114.9 | Very wide | ??? to 77.14 |
| 95% CI, K | ??? | ??? | ??? | 0.02740 to ??? | ??? | ??? | ??? to 0.2722 |
| 95% CI, half-life (min) | ??? | ??? | ??? | ??? to 25.29 | ??? | ??? | 2.546 to ??? |
| 95% CI, Tau (min) | ??? | ??? | ??? | ??? to 36.49 | ??? | ??? | 3.674 to ??? |
| Degrees of freedom | 3 | 3 | 9 | 7 | 2 | 1 | 3 |
| R² | 0.007139 | 0.6048 | 0.2233 | 0.5540 | 0.4023 | 0.6115 | 0.8217 |
| Sum of squares | 108.6 | 40.84 | 150.9 | 40.54 | 68.62 | 25.25 | 1285 |
| Sy.x | 6.015 | 3.690 | 4.094 | 2.407 | 5.857 | 5.025 | 20.70 |
| Constraint on K | K > 0 | K > 0 | K > 0 | K > 0 | K > 0 | K > 0 | K > 0 |
| # X values |  |  |  |  |  |  |  |
| # Y values analyzed | 24 | 24 | 24 | 24 | 24 | 20 | 24 |

*Note.* Compound identities are D-3, H-D-Tyr-Thr-Asp-OH; L-3, H-Tyr-Thr-Asp-OH; D-10, H-D-Trp-Thr-Asp-OH; L-10, H-Trp-Thr-Asp-OH; R-11, H-(R)-Tcc-Thr-Asp-OH; and S-11, H-(S)-Tcc-Thr-Asp-OH. Profile-likelihood confidence-interval fields containing “???” are reproduced exactly from the Prism export and should not be interpreted as bounded confidence intervals. The S-11 fit also returned an unstable plateau/span and an extremely long fitted half-life.

**Table S8. Microscale Thermophoresis Parameters and Data.**

| # | Ligand name | FrhA-PBD concentration | Ligand concentration range | n | Excitation power | MST power | *K*d | Reported *K*d confidence | Regression SE | Signal/noise |
| --- | --- | --- | --- | --- | --- | --- | --- | --- | --- | --- |
| 4 | H-Tyr-D-Thr-Asp-OH | 80 nM | 0.0134–440 µM | 3 | 20% | 40% | 3.04 µM | ± 1.11 µM | 1.707 | 8.87 |
| 5 | H-Tyr-allo-Thr-Asp-OH | 80 nM | 0.0134–440 µM | 3 | 20% | 40% | 7.38 µM | ± 2.33 µM | 1.698 | 10.5 |
| 6 | H-Tyr-D-allo-Thr-Asp-OH | 80 nM | 0.0134–440 µM | 3 | 20% | 40% | 32.4 µM | ± 14.3 µM | 1.657 | 8.79 |
| 7 | H-D-Tyr-allo-Thr-Asp-OH | 80 nM | 0.0134–440 µM | 3 | 20% | 40% | 1.28 µM | ± 0.477 µM | 1.602 | 8.55 |
| 8 | H-D-Tyr-D-Thr-Asp-OH | 80 nM | 0.0134–440 µM | 3 | 20% | 40% | 4.63 µM | ± 1.83 µM | 1.269 | 8.23 |
| 9 | H-D-Tyr-Hyp-Asp-OH | 80 nM | 0.0134–440 µM | 2 | 60% | 40% | No binding† | — | 4.947 | 0 |
| L-10 | H-Trp-Thr-Asp-OH | 20 nM | 0.000977–32 µM | 3 | 20% | 40% | 244 nM | ± 116 nM | 3.299 | 6.91 |
| D-10 | H-D-Trp-Thr-Asp-OH | 20 nM | 0.000977–32 µM | 3 | 20% | 40% | 110 nM | ± 43.5 nM | 1.987 | 8.53 |
| R-11 | H-(R)-Tcc-Thr-Asp-OH | 20 nM | 0.000420–13.8 µM | 3 | 20% | 40% | 102 nM | ± 30.2 nM | 1.128 | 11.5 |
| S-11 | H-(S)-Tcc-Thr-Asp-OH | 80 nM | 0.0134–440 µM | 3 | 20% | 40% | 374 nM | ± 153 nM | 1.319 | 8.78 |
